# Intracellular sterol sensing controls intestinal B cell differentiation

**DOI:** 10.1101/2020.11.03.367177

**Authors:** Bruno C. Trindade, Simona Ceglia, Alyssa Berthelette, Fiona Raso, Kelsey Howley, Jagan R. Muppidi, Andrea Reboldi

**Affiliations:** Department of Pathology, University of Massachusetts Medical School, Worcester, MA, USA; Lymphoid Malignancies Branch, Center for Cancer Research, National Cancer Institute, National Institutes of Health, Bethesda, MD, USA

## Abstract

Intestinal B cell responses are critical for the maintenance of gut homeostasis, yet environmental signals that control B cell metabolism and effector function remain poorly characterized. Here, we show that Peyer’s patches germinal center (GC) B cells are sensitive to 25-hydroxycholesterol (25-HC), an oxidized metabolite of cholesterol produced by the enzyme cholesterol 25-hydroxylase (CH25H). In mice lacking CH25H, antigen-specific GC B cells show an increased cholesterol metabolic signature and preferentially differentiate in plasma cells (PCs), thereby inducing a stronger intestinal IgA response upon immunization or infection. GC B cells express the sterol sensor SREBP2 and use it to sense 25-HC. Deletion of SREBP2 from GC B cells prevents PC differentiation and forces the maintenance of GC identity. GC localized oxysterol production by follicular dendritic cells is central in dictating GC metabolism and imposing B cell fate. Our findings show that the 25-HC-SREBP2 axis shapes B cell effector function in intestinal lymphoid organs and indicate that dietary cholesterol can instruct local B cell response.

## Introduction

In the small intestine, most of the T cell-dependent IgA response originates in Peyer’s patches (PPs), the lymphoid organs closest to the intestinal lumen. The anatomical position of PPs place them at the primary site of absorption for both dietary and bacterial products, therefore exposing them to a variety of metabolic cues. While metabolites absorbed from the intestinal lumen can shape PP organogenesis (van de Pavert et al., 2014) and migration of adaptive immune cells (Mora et al., 2006), mainly via retinoic acid receptor signaling, a precise, mechanistic understanding of how diet-derived metabolites impact B cell differentiation and IgA production in PPs is lacking.

In humans and mice, cholesterol absorption is restricted to the small intestine and cholesterol byproducts (Cyster et al., 2014) possess known modulatory activities on lymphocytes. Diets high in cholesterol are shown to impact adaptive immune responses, including B cell activity (Petta et al., 2018). Among the cholesterol byproducts with immunomodulatory properties, 25-hydroxycholesterol (25-HC) plays a multifaceted role in immune responses. 25-HC, the oxysterol produced by the enzyme cholesterol 25-hydroxylase (CH25H), is the substrate necessary for the generation of 7α,25-HC, the strongest ligand for the orphan G protein-coupled receptor 183 (GPR183, also known as EBI2). Despite 7α,25-HC and 25-HC differing only in the hydroxylated group at the 7α position, 25-HC does not bind and activate EBI2. EBI2 drives migration of several immune cell types (Kelly et al., 2011; Pereira et al., 2009; Yi et al., 2012) (Baptista et al., 2019) (Emgård et al., 2018) (Chu et al., 2018) (Melo-Gonzalez et al., 2019) and CH25H is required for this process. 25-HC can also work in a EBI2-independent role, primarily as an innate immune modulator (Reboldi et al., 2014) (Blanc et al., 2012) (Gold et al., 2014; Liu et al., 2012). (Dang et al., 2017). Whether 25-HC is also involved in adaptive immune responses per se remains to be determined.

25-HC was originally identified as a potent inhibitor of sterol biosynthesis, with higher suppressive potency than cholesterol itself (Goldstein et al., 2006). 25-HC prevents sterol biosynthesis by maintaining inactive sterol response element‐binding proteins 2 (SREBP2) in the endoplasmic reticulum (ER) (Goldstein et al., 2006). When the intracellular level of 25-HC decreases, SREBP2 moves into the Golgi, where it is cleaved by site 1 and site 2 proteases and subsequently activated as a transcription factor (Sakai et al., 1996). Despite its critical role in maintaining cellular cholesterol homeostasis, the role of the SREBP2 pathway in immune cells has been poorly characterized (Kidani et al., 2013), and its impact on B cell fate, especially during intestinal responses, is unknown.

B cell response in the PPs, and the subsequent IgA production, are driven by continual exposure to microbial-derived molecules (Jung et al., 2010) (Reboldi et al., 2016) (Belkaid and Naik, 2013) (Hooper and Macpherson, 2010) as seen by the presence of chronic germinal center (GC) composed of discrete clusters of GC B cells, T follicular helper cells (Tfh cells) and follicular dendritic cells (FDCs) in the center of B cell follicles. PP GC B cells give rise to plasma cells (PCs) and memory B cells (MBCs): however, the impact of intestinal-derived metabolites on GC B cell differentiation remains largely unexplored. In this study, we sought to further define how 25-HC controls intestinal IgA response and to understand how dietary cholesterol restrains PC generation during enteric B cell response.

Here, we report that B cell differentiation to IgA PCs in intestinal lymphoid organs is controlled by the oxysterol 25-HC independently of the GPR183/EBI2 pathway. Mice lacking *Ch25h* have normal IgA class switch recombination *in vivo* but show increased antigen (Ag)-specific IgA response to toxin and enteric pathogen. In PPs, FDCs are required for the generation of the 25-HC niche that mediates suppression of cholesterol metabolic genes in GC B cells. Mechanistically, 25HC inhibits SREBP2 activation, nuclear translocation and transcriptional activity: in the absence of 25-HC, SREBP2 target genes are upregulated in GC B cells and ectopic expression of *Srebf2* in GC B cells drives increased PC differentiation.

In contrast, GC B cells lacking SREBP2 maintain a GC phenotype and have reduced propensity to become PCs both *in vitro* and *in vivo*. Our results establish a role for a dietary-derived sterol in controlling B cell fate in the mucosal draining lymph node during T cell-dependent IgA response.

## Results

### CH25H controls antigen-specific intestinal IgA response independently of EBI2

To measure the impact of the 25-HC producing enzyme CH25H on mucosal B cell responses, we immunized *Ch25h*^*-/-*^ mice and co-housed, littermate controls (LMCs) with the intestinal antigen cholera toxin (CT) and quantified the IgA response in the intestinal lamina propria (LP). While the IgA^+^ PC fraction (**Fig**.**1A**) was identical between genotypes, CT-specific IgAsecreting cells (CT-IgA-ASC) were present in higher numbers in the duodenum of mice lacking CH25H (**Fig**.**S1A**): the total number of IgA-ASC was not different between genotypes (**Fig**.**S1B**), leading to a higher frequency of CT-IgA-ASC in *Ch25h*^*-/-*^ mice (**Fig**.**1B**). In line with the overall distribution of IgA-ASC in the small intestine (**Fig**.**S1C**), we only observed a consistent frequency of IgA-ASC in the duodenum, with very little IgA or CT-IgA-ASC in the ileum of *Ch25h*^*/-*^ mice (**Fig**.**S1D**,**E**). Since CH25H is required for the generation of 7a,25-HC, the most potent ligand for EBI2, a GPCR that is expressed on B cells and controls B cell positioning and differentiation, we tested whether CH25H deficiency might impact Ag-specific IgA response through EBI2 by immunizing *Ebi2*^*-/-*^ and LMCs. *Ebi2*^*-/-*^ mice showed unchanged number of CT-IgA-ASC, IgA-ASC (**Fig**.**S1F, G**), and overall frequency of CT-IgA-ASC (**Fig**.**1C**) suggesting that CH25H controls the intestinal humoral response independently of EBI2 migration. CT-specific IgA titers were increased in the small intestinal lavage (**Fig**.**1D**), but not in the serum (**Fig**.**1E**), highlighting that CH25H is predominantly affecting intestinal, but not systemic, antigen-specific IgA responses.

**Figure 1.**
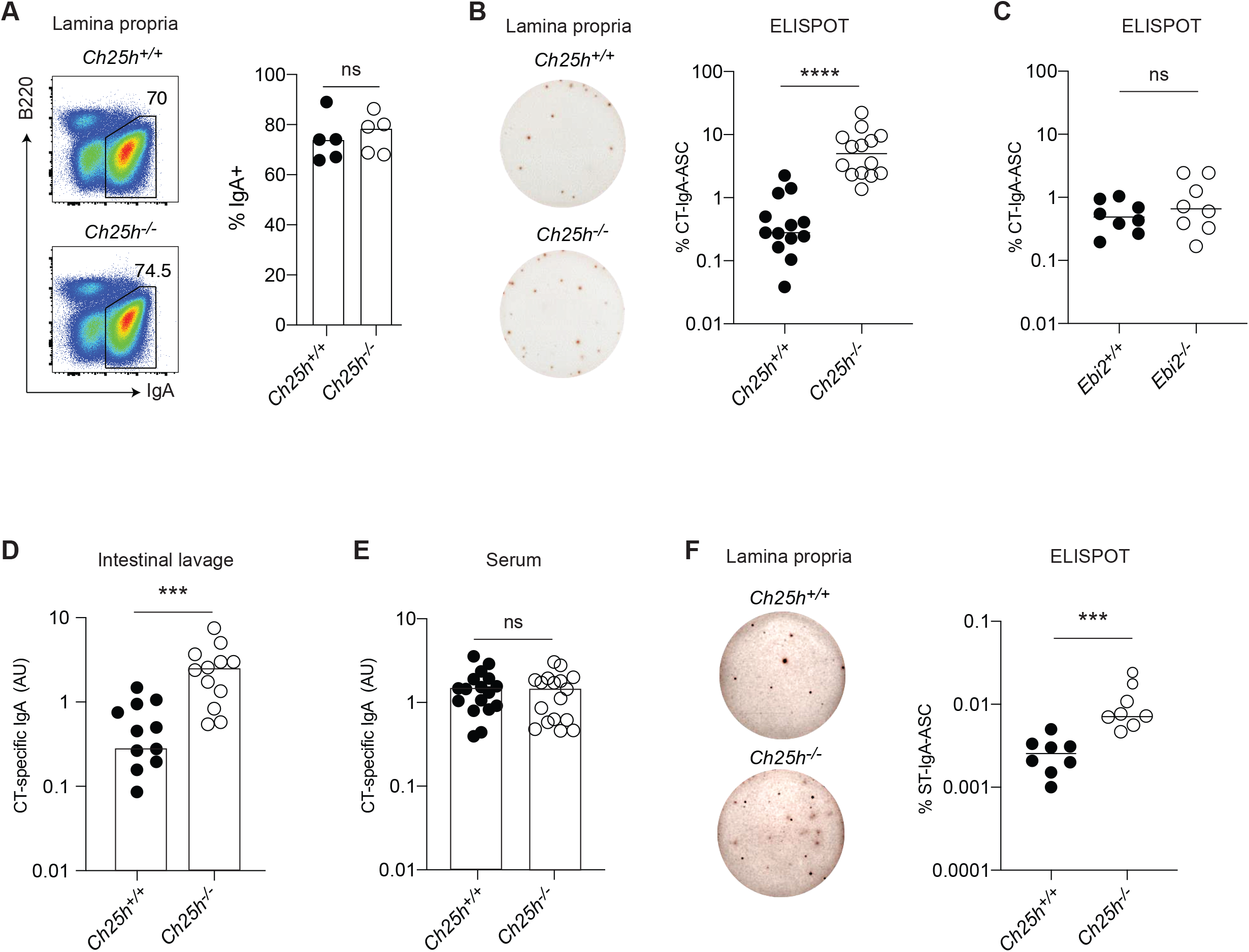
25-HC, but not EBI2, shapes antigen-specific IgA response in duodenal LP. (A) Representative Flow cytometric analysis of IgA+ plasma cells in Duodenum (LP) of *Ch25h*^*-/-*^ and LMC mice. (B) ELISPOT of isolated lamina propria PC from *Ch25h*^*-/-*^ and (C) *Ebi2*^*-/-*^ with quantified percentage of CT-specific IgA secreting PC. (D,E) Quantification of secreted CT-IgA by ELISA assay in small intestine lavage and serum of mice immunized for 3 weeks with CT. (F) Percentage of Salmonella T.-specific IgA PC in intestinal lamina propria of *Ch25h*^*-/-*^ and LMC mice, analyzed by ELISPOT assay two weeks post-infection. Data are representative of three independent experiment, ns=non-significant, *** p<0.005; ****p<0.001 (unpaired Student’s T test).

To further investigate the role of CH25H on intestinal IgA response, we infected *Ch25h*^*-/-*^ mice and LMCs with *aroA*deficient (D*AroA*) *Salmonella enterica Serovar Typhimurium* (*ST*), a defective strain of ST that has an intact route of entry into PPs but reduced infectivity (Martinoli et al., 2007), allowing for the measurement of IgA response in B6 mice (Monack et al., 2004). In absence of CH25H, we observed an increased number of ST-IgA-ASC in the LP (**Fig**.**1F**).

Overall, our results showed that mice lacking CH25H are characterized by an enhanced Ag-specific IgA response against toxin and enteric bacteria, suggesting that 25-HC can control T cell-dependent IgA responses.

### PC-generation, not class switch recombination, is controlled by 25-HC in PPs

IgA-secreting PCs represent the terminal differentiation step of B cells stimulated by intestinal antigen in the PPs. Upon Ag stimulation and GC entry, GC B cells differentiate to IgA PC that egress from the PP and migrate to LP using a combination of integrin and chemokine receptors. We therefore asked whether CH25H’s effect on Ag-specific IgA ASC generation was anatomically restricted and initiated in PPs. To this end we took advantage of FTY720, a sphingosine 1-phosphate receptor (S1PR1) antagonist that can prevent IgA PC egress from PPs (Gohda et al., 2008).

We immunized *Ch25h*^*-/-*^ mice and LMCs with CT and administered FTY720 to trap newly generated, Ag-specific PCs in PPs. As expected, FTY720 treatment increased the number of IgA-ASC in the PPs (**Fig**.**2A**, **Fig**.**S2A**), but only *Ch25h*– deficient mice showed an enhanced Ag-specific PC response (**Fig**.**2B, Fig**.**S2B**). In contrast, mice lacking EBI2 showed no increase in CT-response, highlighting how CH25H, but not EBI2, is involved in controlling specific IgA response in PPs (**Fig**.**2C**, **Fig**.**S2C**). In line with the data from LP, blocking of S1PR1-dependent ASC egress with FTY720 during ST infection also enhanced IgA-ASC response in PPs of *Ch25h*^*-/-*^ mice. (**Fig**.**2D**). While *Ebi2*^*-/-*^ mice have showed systemic decrease of PC response, we did not observe an effect on PC generation in PPs (**Fig**.**S2D**), and *Ch25h*^*-/-*^ mice, but not *Ebi2*^*/-*^ mice, were able to generate more Ag-specific PCs. These results suggest the existence of a process dependent on CH25H, but independent of EBI2, that shapes IgA specific response in PPs.

Previous work has suggested that 25-HC could block IgA class switching (Bauman et al., 2009) however ex-vivo FACS analysis of B cell subsets from PPs and mesenteric LN (mLN) in *Ch25h*^*+/-*^ and *Ch25h*^*-/-*^ showed that lack of 25-HC has no impact on the ability of B cells to undergo IgA class switch recombination (**Fig**.**2E**, **Fig**.**S2E**).

**Figure 2.**
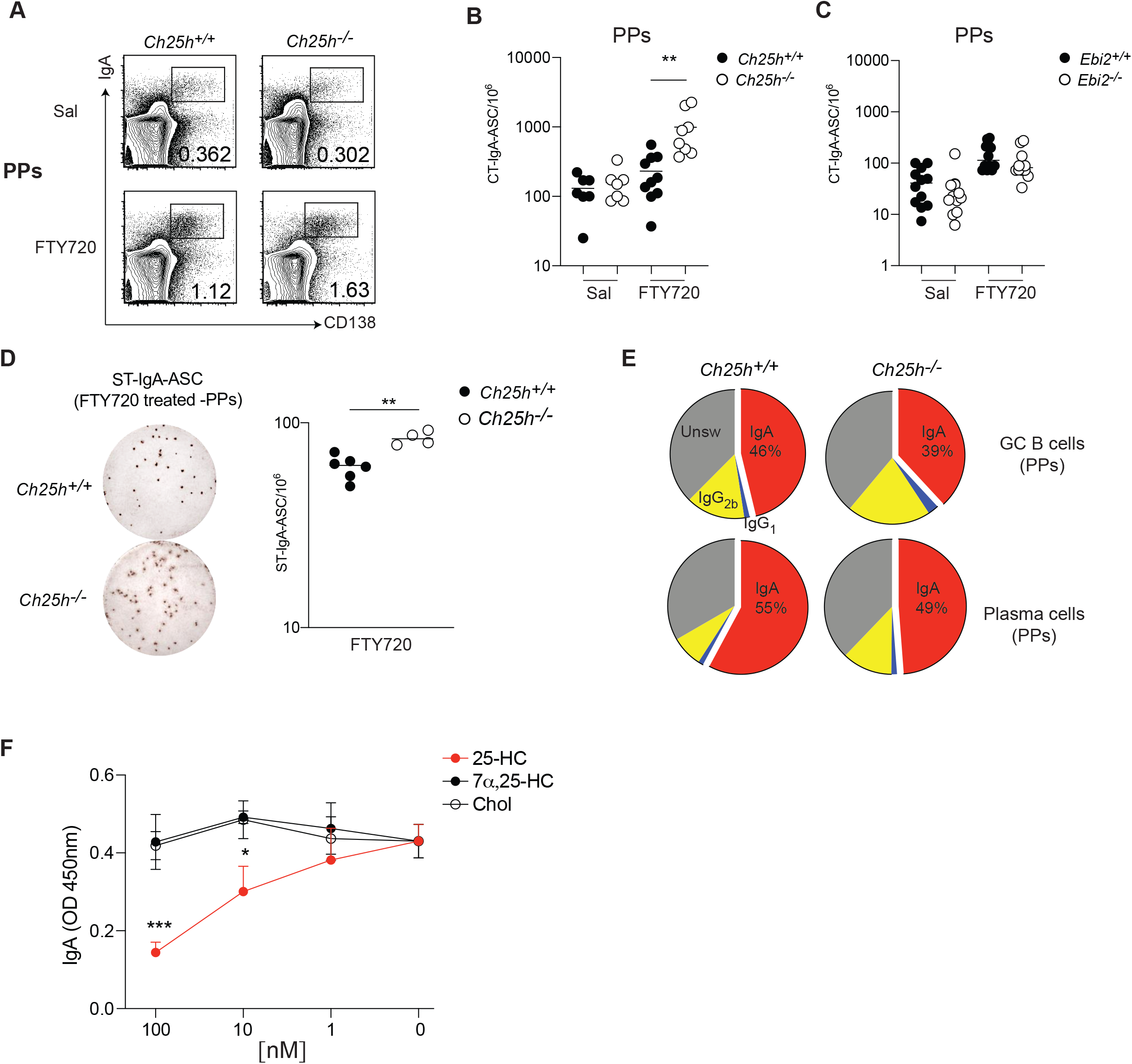
25-HC modulates Ag-specific PC generation in PPs. (A) Flow cytometric analysis of plasma cells in PP of *Ch25h*^*-/-*^ and LMC mice treated daily with FTY720 or saline for one week. (B) Number of CT-specific IgA secreting cells in *Ch25h*^*-/-*^,(C) *Ebi2*^*-/-*^ and wild type mice, measured by ELISPOT. (D) Representative ELISPOT showing Salmonella T. specific IgA secreting cells in PP of *Ch25h*^*-/-*^ and LMC mice treated with FTY720. Data indicate the frequency of ST-ASC among PP cells. (E) Analysis of Class switching recombination in PP by Flow cytometry. (F) Quantification of secreted IgA by ELISA from GC B cells cultured with NB21 cells and incubated with the indicated sterols for 3.5 days. A,B,C,E from at least three independent experiments, D, F from two independent experiments. Statistical significance was measured by two-way ANOVA using Bonferroni’s corrections in B, C, F **p<0.01 (two-way ANOVA using Bonferroni’s corrections (B, F) unpaired Student’s t test (D))

To assess whether 25-HC could directly restrain PC generation from GC B cells, we used NB21 culture assay, in which the feeder cell layer expresses CD40L, BAFF and IL-21 (Kuraoka et al., 2016). Sorted GC B cells were cocultured with NB21 and then treated with different oxysterols: 25-HC, but not 7a,25-HC or cholesterol, decreased the Ab-secreting cell generation (**Fig**.**2F**). Similar results were obtained with GC B cells stimulated with anti-CD40 and cultured with NB21 supernatant, ruling out a direct effect of 25-HC on NB21 cells (**Fig**.**S2F**), and suggests that 25-HC can act directly on GC B cells and restrain their ability to differentiate in PCs. Overall, our data suggest that intestinal and systemic antibody responses are differentially dependent on oxysterols, with 25-HC controlling local Ag-specific PC response in an EBI2independent fashion during enteric response.

### FDCs contribute to 25-HC PP levels and dietary cholesterol reduces IgA response via CH25H

25-HC can be secreted (Reboldi et al., 2014) and diffuse to the surrounding tissue (Cyster et al., 2014; Yi et al., 2012) to generate a 25-HC niche, a microanatomical area rich in 25-HC that can regulate nearby immune cells in a EBI2-independent way. *Ch25h* expression has been reported in several cell types of both hematopoietic and non-hematopoietic origin, mainly in the context of EBI2 ligand generation: however, the contribution of *Ch25h-*expressing cells to the 25-HC niche *in vivo* is less clear. While the EBI2 ligand 7a,25-HC can be easily measured in a transwell assay based on EBI2-mediated migration, 25-HC does not bind EBI2 and therefore remains undetected in that setting. To unambiguously measure 25-HC in tissue extracts, we designed a modified transwell assay and verified it using synthetic oxysterols (**Fig**.**S3A**). We first inactivated 7α,25-HC by incubating it with 293T cells expressing HSD3B7, a 7a-oxidized cholesterol dehydrogenase enzyme that oxidizes all existing EBI2 ligand; subsequently we harvested and incubated 25-HC-containing extract with 293T cells expressing CYP7B1. This second incubation converted 25-HC to 7α,25-HC, which was now measured in the transwell using EBI2-expressing M12 cells (**Fig**.**S3B**). Analysis of intestine and intestinal secondary lymphoid organs (SLO) showed that 25-HC can be measured in all of these tissues, and requires the activity of CH25H enzyme: thus, intestinal cells producing 25-HC that is not converted into EBI2 ligand, exist *in vivo* in unmanipulated mice (**Fig**.**S3C**).

PC differentiation largely takes place in the GC, so we reasoned that a 25-HC niche anatomically restricted to the GC could influence activated B cell fate decision. In GC, Follicular dendritic cells (FDCs), stromal cells (SCs) that display antigen in the light zone for BCR testing, have high levels of *Hsd3b7* transcripts (Yi et al., 2012) and can inactivate EBI2 ligand, thereby maintaining EBI2 expressing follicular B cells, outside of the GC, while allowing GC B cells, which do not express EBI2, to reach and remain inside the GC structure (Cyster et al., 2014). As 25-HC is not sensitive to HSD3B7 activity, the GC environment is likely to contain higher concentrations of 25-HC compared to the follicles: since EBI2 ligand regulates naïve B cell positioning, we posit that most of 25-HC in the follicles is not freely available as it is converted to 7a,25-HC by CYP7B1 (**Fig**.**S3D**). FDCs express *Ch25h* (Rodda et al., 2018), but whether FDCs are critical to establish GC 25-HC is unknown.

We employed two different strategies to assess whether FDCs are the primary source of 25-HC in PPs. First, we used CD21DTR mice, where *Cd21*-Cre mice are crossed with mice carrying simian diphtheria toxin receptor (DTR) downstream of a floxed stop element in the ROSA26 locus. We generated bone marrow (BM) chimeras using CD21-DTR mice (or LMCs) as hosts: in this setting the DTR is expressed only in non-hematopoietic CD21-expressing cells, i.e. FDCs (Wang et al., 2011). Short term DT injection led to FDC ablation: while we only observed a small reduction in EBI2, 25-HC was largely reduced in PPs (**Fig**.**3A**), suggesting that cells other than FDCs might contribute to the follicular production of EBI2 ligand (Rodda et al., 2018). As a second test of the FDC requirement in 25-HC niche generation, we depleted FDCs by blocking lymphotoxin (LT) signaling through LT-beta receptor (LTbR) on FDCs using soluble LTbR fused to human IgG Fc (LTbRFc). The blockade prevents the interaction between lymphotoxin-a_2_b_1_-expressing GC B cells and LTbR-expressing FDCs, leading to FDC death (Mackay et al., 1997). FDC depletion only marginally impacted EBI2 ligand generation but led to a dramatic reduction of 25-HC (**Fig**.**3B**), strongly suggesting that FDCs primarily contribute to the 25-HC niche in the GC. Since CH25H requires cholesterol to generate 25-HC we asked whether changes in dietary cholesterol content could shape the intestinal 25-HC niche. We fed mice with normal food (NF) or normal food with the exclusive addition of 0.15% cholesterol found in the Western diet (high cholesterol food, HCF) for 1 week and measured 25-HC in PPs. We found that 25-HC was increased in animal fed HCF (**Fig**.**3C**), suggesting that altering the intestinal absorption of dietary cholesterol can shape the 25-HC niche. EBI2 ligand was also increased in PPs, highlighting a previously unreported link between dietary cholesterol and immunomodulatory sterols.

We then asked whether diet-dependent changes in the PP 25-HC niche could influence B cell fate: mice fed with different diets were immunized with CT and treated with FTY720. WT mice fed HCF showed decreased generation of CT-IgA-ASC compared to WT mice fed with NF. In contrast, *Ch25h*^*-/-*^ mice showed only a modest reduction in CT-IgA-ASC (**Fig**.**3D**). We were able to observe a similar effect of HCF and CH25H deficiency on CT-IgA-ASC in the mLN (**Fig**.**S3E**,**F**), suggesting that 25-HC dependent restraint on PC generation occurs in other intestinal SLOs.

**Figure 3.**
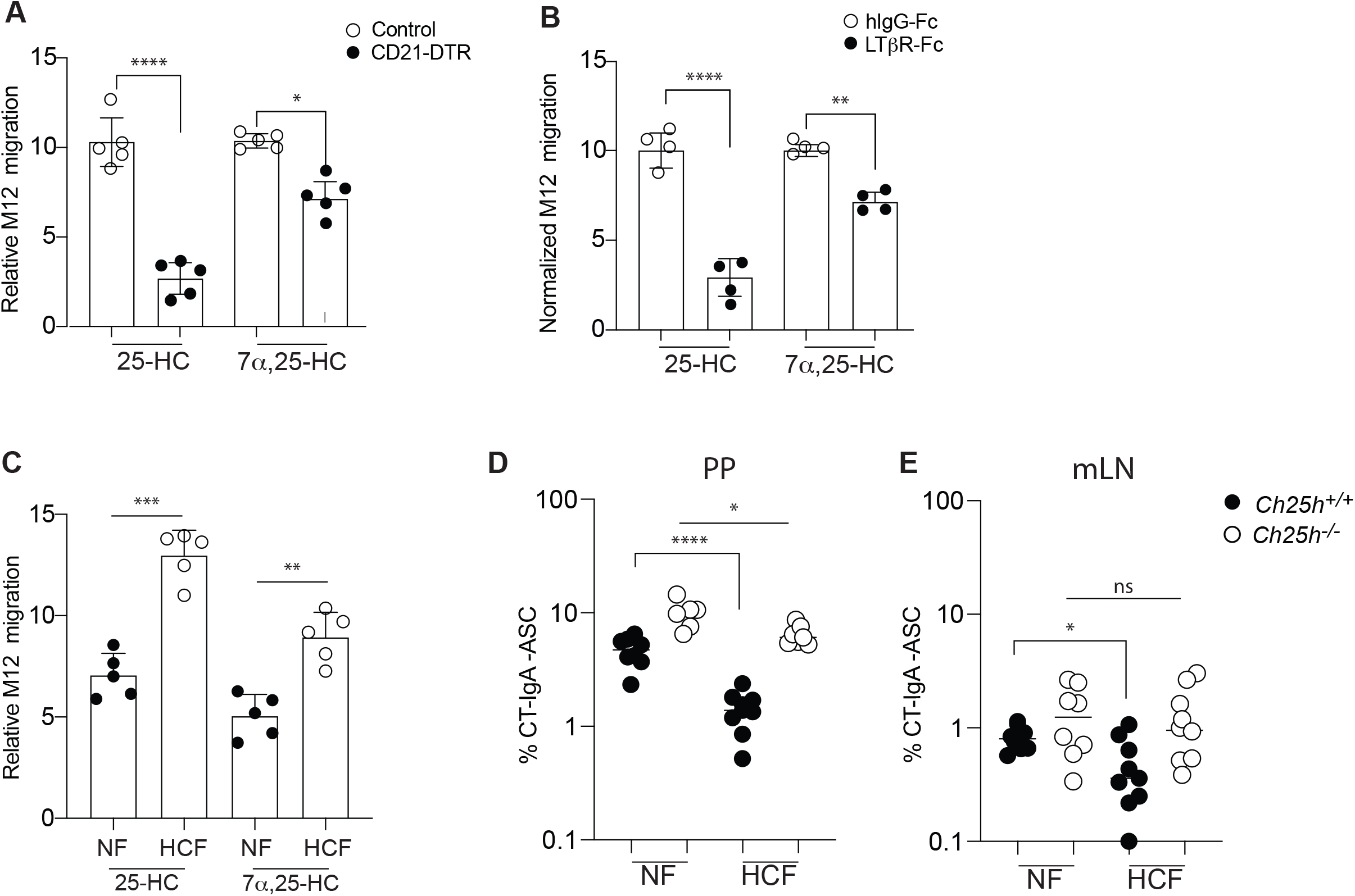
FDCs and diet shape tissue 25-HC levels and control IgA response. (A) CD21-DTR and LMC mice were reconstituted with WT bone marrow and treated with diphtheria toxin for 16h: lipids were extracted from PPs for 25-HC measurement using transwell migration assay with EBI2^+^ M12 cells. (B) Transwell migration assay of EBI2^+^ M12 cells in response to lipid extracted from PP of wild type mice treated with recombinant LTbR-FC or isotype control. (C) Relative migration of EBI2^+^ M12 cells exposed to PP lipid extracts of WT mice treated for 1 weeks with 0.15 % High cholesterol diet (Western diet) or normal chow. (D,E) Frequency of CT-specific ASC detected by ELISPOT of mLN and PP suspension cells from *Ch25h*^*-/-*^ and LMC fed with mice normal chow or 0.15% of HCF. Results are pooled from 3 or 4 independent experiments. ns=non-significant, *p<0.05, **p<0.01,***p<0.005, ****p<0.001 (twoway ANOVA)

### SREBP2 activation is induced by BCR signaling and is controlled in vivo by 25-HC

Due to the role of 25-HC in controlling activation and nuclear translocation of sterol response element‐binding proteins (SREBP2) (Goldstein et al., 2006), we analyzed the SREBP2 transcriptional activity in B cells. Since SREBP2 regulates its own transcription (Sato et al., 1996), we initially measured *Srebf2* transcript in PP B cell subsets (**Fig**.**4A**). GC B cells and PC showed the highest level of *Srebf2*, followed by MBCs, while follicular B cells showed the lowest *Srebf2* transcription. Similar dynamics of *Srebf2* transcription was observed in B cell subsets from the mLN (**Fig**.**S4A**). Transcripts for *Srebf1a*, an isoform of SREBP1 protein than can control certain aspects of lipid metabolism, were detected at a considerably lower levels in both PPs and the mLN, with both GC and PC transcribing *Srebf1a*. (**Fig**.**4B**, **Fig**.**S4B**). To understand whether 25HC can control SREBP2 levels *in vivo* we analyzed *Srebf2* transcript and SREBP2 target gene transcripts in *Ch25h*^*-/-*^ mice. *Srebf2* expression is increased in GC B cells and PCs in both PPs and mLN in absence of 25-HC (**Fig**.**4C**, **Fig**.**S4C**) while no difference was observed in follicular B cells and a minimal increase was present in MBCs. These results demonstrate that *in vivo* 25-HC levels can control SREBP2 activation in some antigen-experienced B cell subsets.

**Figure 4.**
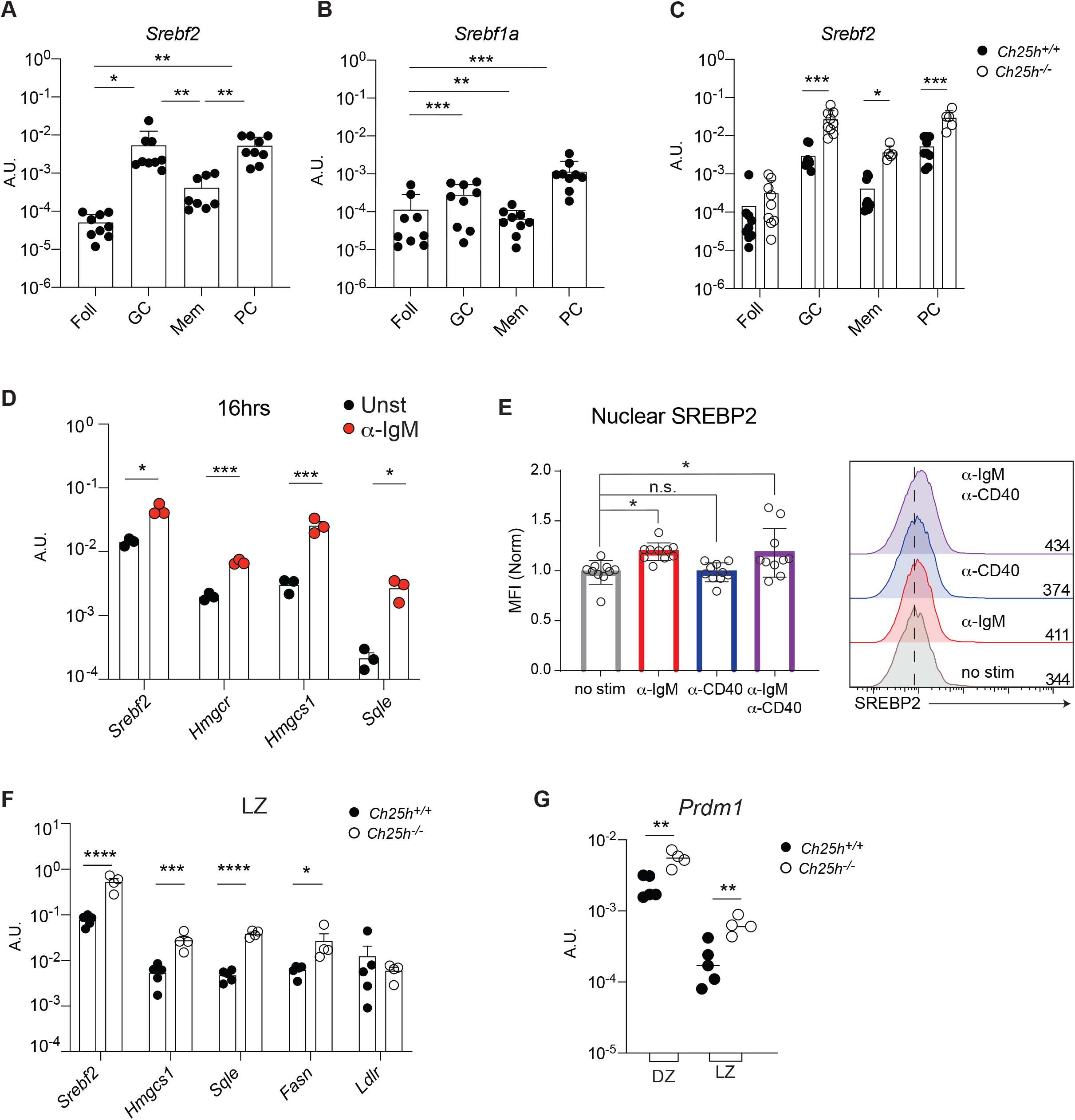
25-HC regulates SREBP2 transcription and activation in activated B cells. Transcript level of *Srebf2* (A) and *Srebf1a* (B) in B cell subsets from PP of wild-type mice and (C) from PP of *Ch25h*^*-/-*^ and LMC mice measured by qPCR. (D) Analysis of gene expression of *Srebf2* and SREBP2-target in follicular B cells stimulated with anti-IgM for 16h. (E) Cumulative and representative Flow cytometry of SREBP2 nuclear staining in Follicular B cells stimulated with the indicated stimuli F) qPCR results of *Srebf2* and its target genes in sorted B cells of germinal center LZ. (G) Blimp expression in sorted DZ (CXCR4+CD86-) and LZ (CXCR4-CD86+) GC B cells from PP of *Ch25h*^*-/-*^ and LMC mice. Each symbol represents one independent mouse from 9-10 (in A,B,C, E) or 4 mice (in F,G). Graphs show mean and SEM. *p<0.05, **p<0.01,***p<0.005, ****p<0.001(two-way ANOVA)

SREBP2 induction is generally driven by low levels of intracellular sterols, including selected oxysterol concentrations, but whether other signals can control SREBP2 activation is unclear. All activated B cell subsets, but not naïve B cells, upregulate *Srebf2* transcripts to some extent, thus we reasoned that BCR stimulation and T cell help could possibly induce SREBP2 activation. Follicular B cells stimulated with anti-IgM for 16h showed *Srebf2* induction and increase in SREBP2 target genes *Hmgcr, Hmgcs1* and *Sqle* (**Fig**.**4D**). In contrast anti-CD40 stimulation, that mimics T cell help, had little effect on SREBP2-target *Hmgcr, Hmgcs1* genes at a later time point compared to stimulation with both anti-IgM and anti-CD40, but it was able to upregulate *Sqle* transcription (**Fig**.**S4D**). Transcription of SREBP2 target genes suggest that SREBP2 abundance is increased in BCR-stimulated B cells. We first analyzed the amount of SREBP2 by staining stimulated B cells intracellularly with an antibody that recognizes a site in SREBP2 that is only present in the ER-resident, unprocessed SREBP2. In line with the qPCR data, BCR stimulation drove SREBP2 upregulation (**Fig**.**S4E**). SREBP2 function is mediated by its nuclear translocation. Therefore we sought to investigate the cellular dynamics of SREBP2 in stimulated B cells by flow cytometry analysis of transcription factor (TF) in isolated cell nuclei (Gallagher et al., 2018). We validated the nuclear isolation by analyzing the B cell line WEHI expressing cytosolic GFP, nuclear RFP (H2B-RFP) and stained with fluorescent ER tracker: this strategy allowed for the quantification of nuclear TFs at single cell level without ER contamination, which is critical since inactive SREBP2 is sequestered in the ER (**Fig**.**S4F**). Using an antibody directed against the N-terminus of SREBP2, that functions as TF, B cells stimulated through the BCR induced SREBP2 activation and its nuclear translocation (**Fig**.**4E**). IRF4, a transcription factor that is upregulated upon BCR stimulation and is critical for PC formation, also translocated to the nucleus in a similar fashion (**Fig**.**S4G**). SREBP2 nuclear translocation in response to BCR stimulation was dependent by Bruton’s tyrosine kinase (BTK), as BTK inhibitor Ibrutinib maintained SREBP2 sequestered in the cytoplasm (**Fig**.**S4H**).

Since SREBP2 induction is controlled by BCR stimulation, and the 25-HC niche requires FDCs, cells responsible for displaying antigen to GC B cells for BCR testing, we hypothesized that modulation of SREBP2 by 25-HC would be restricted to the GC in the light zone (LZ). Sorted LZ and DZ GC B cells from *Ch25h*^*-/-*^ and LMC mice clearly indicated that both *Srebf2* and SREBP2 target genes are induced to a greater extent in LZ compared to DZ and restrained by 25-HCs (**Fig**.**4F**, **Fig**.**S4I**). Since CH25H deficiency leads to increased GC-derived PC, we analyzed *Prdm1*, the gene that encodes for Blimp-1, the master transcription factor required for PC differentiation. We observed that LZ and DZ GC B cells from PPs of *Ch25h*^*-/-*^ mice upregulate *Prdm1* compared to LMCs GC B cells (**Fig**.**4G**).

Together, our data demonstrate that *in vivo*, 25-HC restrains cholesterologenic genes transcription in anatomically distinct subsets of GC B cells and highlights a process for how 25-HC could restrain initiation of PC transcriptional program in GC B cells, possibly via SREBP2 inhibition.

### Ectopic SREBP2 activation is sufficient to drive PC differentiation

To test whether SREBP2 activation could drive B cell differentiation into PCs per se, we sought to manipulate SREBP2 cellular localization, and therefore activity, in PP GC B cells. We retrovirally transduced BM cells expressing Cre in the *Aicda* locus and tdTom in the ROSA26 locus(*Aicda*^*Cre/+*^ *Rosa26* ^*tdTom/+*^) with retrovirus encoding constitutively active nuclear (n) form of SREBP2 downstream cassette of *lox*P site–stop sequence– *lox*P site cassette (followed by an internal ribosomal entry site and a CD90.1 (Thy-1.1) reporter), which results in restricted expression of nSREBP2 to GC B cells and post-GC B cells (MBCs and PCs) (**Fig**.**5A**cd)Overexpression of nSREBP1a, which is not controlled by 25-HC, was used as control. Since nSREBPs can translocate into the nucleus despite high intracellular sterol concentration, this approach uncouples intracellular metabolism and SREBP2 activation in the context of B cell differentiation. Active SREBP2 led to a striking increase in PC generation in PPs, while active SREBP1a did not result in preferential PC fate compared to empty vector (EV) transduced cells (**Fig**.**5B, C**). SREBP1a overexpression reduced GC output, albeit not at the level of SREBP2, possibly through modulation of fatty acid biosynthesis. The increased PC generation in B cells with active nSREBP2 and the unaffected B cell fate in presence of nSREBP1a is in line with the notion that 25-HC specifically controls SREBP2, but not SREBP1 processing (Reboldi et al., 2014). Analysis of LP PCs, identified by the surface amino acid transporter CD98 (Tellier et a. 2016)(**Fig**.**5D)**, revealed that transduced B cells were able to reach the tissue (**Fig**.**5E**). However, comparison of transduced cell frequency in PP and LP revealed that SREBP-2 transduced cells were present in higher frequency compared to both EV and SREBP1a transduced cells, showing that increased SREBP2 activity is more efficient in generating LP-homing PCs (**Fig**.**5F**). These data establish that alteration of SREBP2 activity instruct B cells to assume distinct fates.

**Figure 5.**
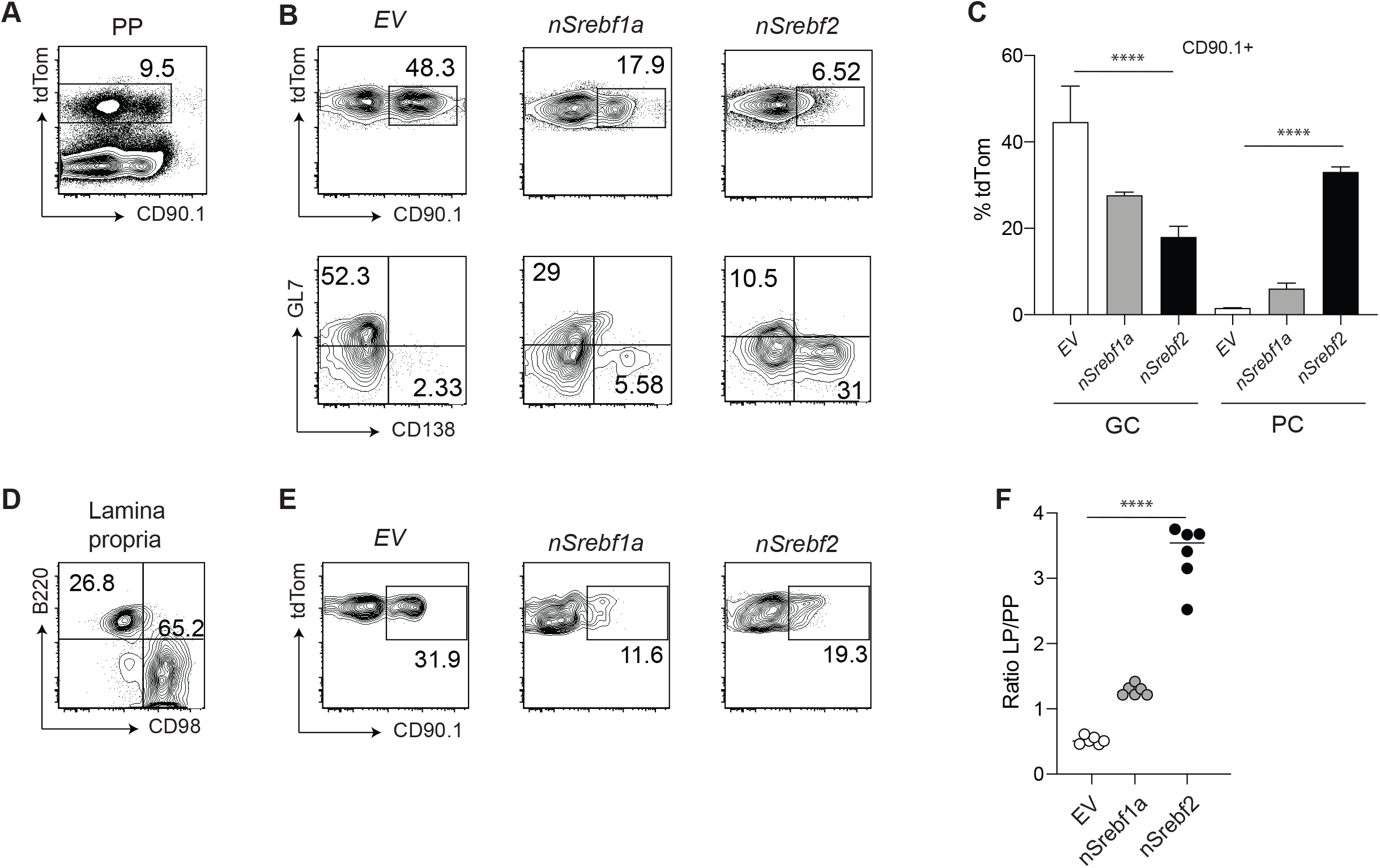
SREBP2 overexpression induces PC differentiation. (A) Representative plot of *Aicda*^*Cre/+*^ *Rosa26* ^*tdTom/+*^ BM transduced with CD90.1 retroviral vector in PPs. (B) Representative plot of frequency and fate of tdTom+ B cells transduced with indicated vectors (C) Summary of the data in (B) (D) Representative gating of tdTom+ cells in lamina propria. (E) Representative plot of frequency and fate of tdTom+ B cells transduced with indicated vectors. (F) Ratio of transduced B cells in PP and LP from data in (B) and (E). Data represents 3 independent experiments. Graphs show mean and SEM. ****p<0.001(two-way ANOVA).

### SREBP2 is required for the generation of PC in vitro and in vivo

To assess whether SREBP2 is necessary for PC differentiation, we generated *Aicda*^cre^ *Srebf2* ^*flox/flox*^ mice: in these animals GC B cells are unable to transcribe and activate SREBP2. We sorted GC from *Aicda*^cre^ *Srebf2* ^*flox/flox*^, *Aicda*^cre^ *Srebf2* ^*flox/+*^ and *Aicda*^cre^ *Srebf2* ^*+/+*^ and assessed their potential for PC differentiation by co-culturing them with NB21 cells: *Srebf2*deficient GC B cells were largely unable to differentiate into PCs and maintained their GC B cell identity (**Fig**.**6A**,**B**,**C**). In line with previous reports of SREBP2 transcriptional activity, *Srebf2* gene dosage can affect cell fate, with GC B cells from *Aicda*^cre^ *Srebf2* ^*flox/+*^ showing an intermediate PC output compared to GC B cells from *Aicda*^cre^ *Srebf2* ^*flox/flox*^ and *Aicda*^cre^ *Srebf2* ^*+/+*^ (**Fig**.**6A**,**B**,**C**). Overall cellular recovery was reduced in cultures containing GC B cells from *Aicda*^cre^ *Srebf2* ^*flox/flox*^, suggesting that SREBP2 deficiency was impairing transition to PCs and impacting GC survival (**Fig**.**S5A**). IgA secretion was also reduced from *Aicda*^cre^ *Srebf2* ^*flox/flox*^ GC B cell cultures (**Fig**.**6D**): since IgA class switch was unaffected in GC B cells lacking SREBP2 (**Fig. S5B**), our data show that SREBP2 is required for allowing efficient GC differentiation into PCs. While *Aicda*^cre^ *Srebf2* ^*flox/flox*^ did not show altered overall IgA-ASC in LP (**Fig**.**S5C**), oral immunization with CT led to a drastic decrease of Ag-specific-IgA ASC and titer (**Fig**.**6E**,**F**).

To finally assess the role of SREBP2 in GCs during intestinal response, we infected and *Aicda*^cre^ *Srebf2* ^*+/+*^ and *Aicda*^cre^ *Srebf2* ^*flox/flox*^ mice with D*AroA ST* and administered FTY720. We observed a decreased number of ST-IgA-ASC in both PPs and mLN in absence of SREBP2 in GC B cells (**Fig**.**7A**, **S6A**) and FACS analysis revealed a cleared reduction in IgA+ PCs (**Fig**.**7B**, **S6B**). In line with in vitro experiments described above, removal of *Srebf2* in GC B cells interfered with normal B cell output, leading to an overrepresentation of GC B cells at the expense of PCs (**Fig**.**7C**,**D**; **S6C**,**D**).

Taken together, our findings show an undescribed role for the sterol sensor SREBP2 in controlling B cell fate during intestinal immune response.

**Figure 6:**
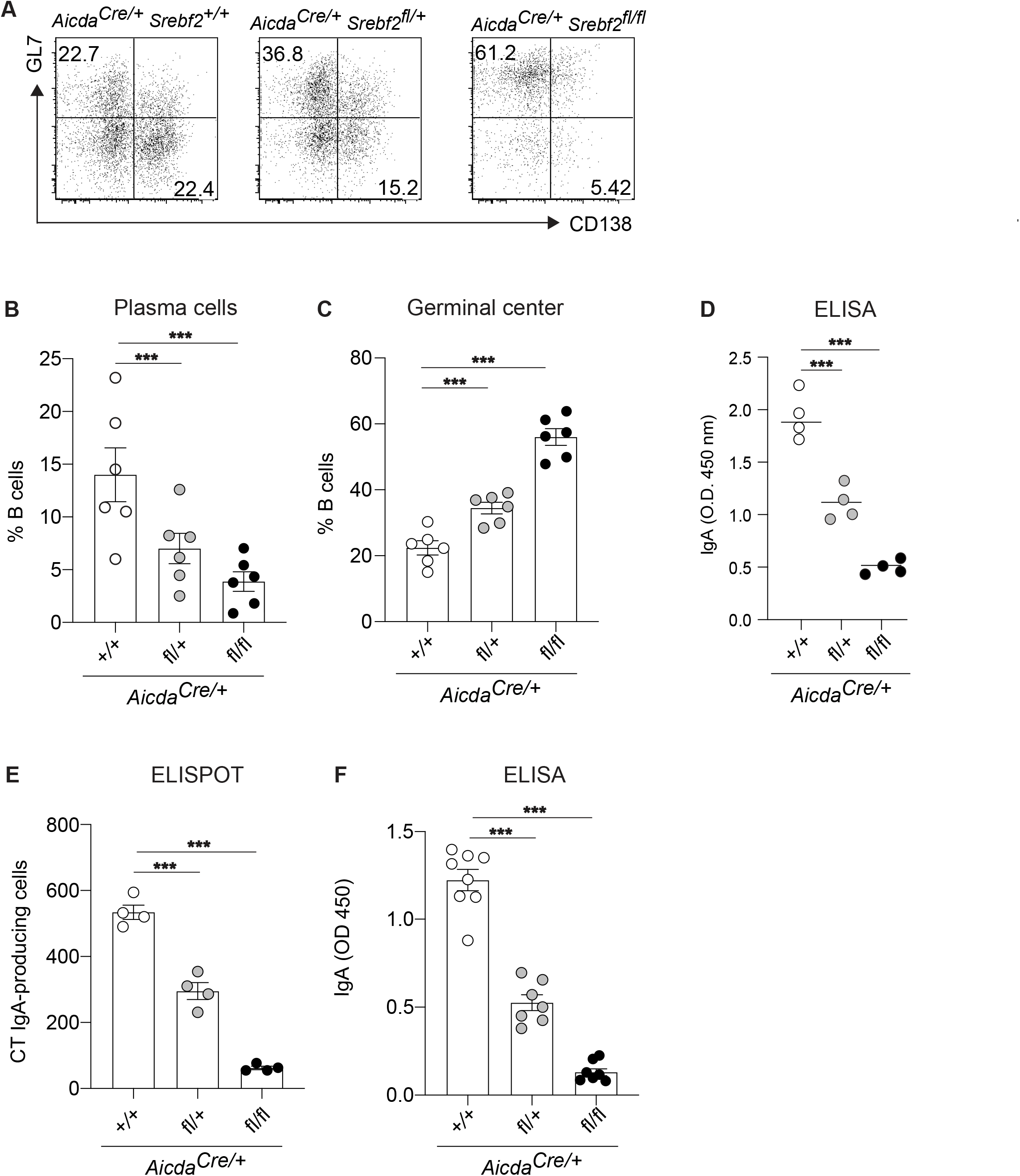
SREBP2 controls GC differentiation potential in vitro and in vivo. (A) Representative flow cytometry of GC B cells from mice of the indicated genotype cultured with NB21 cells for 2.5 days. (B,C) Frequency of GC B cell and PC from A. (D) Quantification of secreted total IgA in A. (E) Number of CT-IgA ASC cells in LP of *Aicda*^cre^ *Srebf2* ^*flox/flox*^ and LMC mice, quantified by ELISPOT. (F) Quantification of CT-IgA in small intestine lavage of mice treated 3 weeks with CT. Each symbol represents one mouse from 2-3 independent experiments. ***p<0.005 (two-way ANOVA).

**Figure 7:**
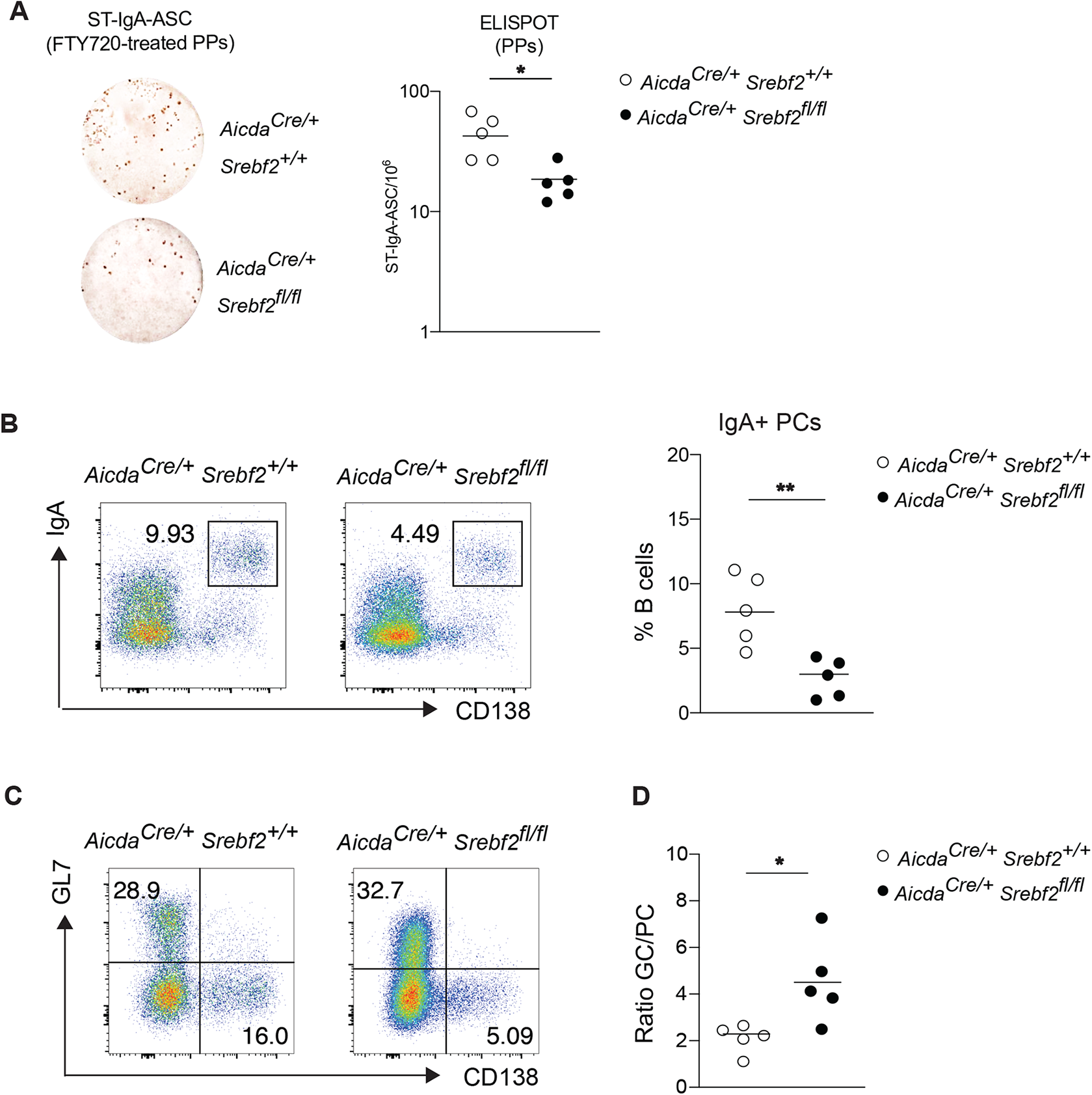
SREBP2 controls intestinal B cell response during enteric infection. (A) Representative ELISPOT and compiled data showing Salmonella-specific IgA secreting cells in PPs of *Aicda*^cre^ *Srebf2* ^*+/+*^ and *Aicda*^cre^ *Srebf2* ^*flox/flox*^ mice infected with Salmonella and treated with FTY720. (B) Representative flow cytometry and frequency of IgA+ PCs from PPs of *Aicda*^cre^ *Srebf2* ^*+/+*^ and *Aicda*^cre^ *Srebf2* ^*flox/flox*^ mice infected with Salmonella and treated with FTY720 as in A. (C) Representative flow cytometry of GC B cell and PC from PPs of *Aicda*^cre^ *Srebf2* ^*+/+*^ and *Aicda*^cre^ *Srebf2* ^*flox/flox*^ mice infected with Salmonella and treated with FTY720 as in A, B. (D) Ratio of GC B cell and PC from PPs from C. Each symbol represents one mouse from 2 independent experiments. *p<0.05, **p<0.01 (two-way ANOVA).

## Discussion

In summary we find that oxysterol 25-hydroxycholesterol (25-HC) acts on GC B cells in PPs to restrain PC differentiation via inhibition of SREBP2 activation. Our data support a novel, direct effect of 25-HC on adaptive immune cells independent of its role in the generation of 7α,25-dihydroxycholesterol (7α,25-HC), the ligand for the surface receptor EBI2.

We have provided evidence that in absence of 25-HC-producing enzyme CH25H, the Ag-specific IgA PC response is enhanced during oral immunization and enteric infection. CH25H is required in PPs to control PC output during T celldependent responses, without any measurable effect on IgA CSR. Our *in vitro* experiments suggest that 25-HC directly affects the ability of GC B cells to undergo PC differentiation. Our data also showed that FDCs are the dominant cell type required to establish the 25-HC niche in the GCs. FDCs have a well-established role in maintaining the anatomical organization of follicular and GC response by enzymatic degradation of metabolites acting on GPCR as described for the EBI2 ligand 7α,25-HC (Yi et al., 2012) and the P2YR8 ligand S-geranylgeranyl-L-glutathione (Lu et al., 2019; Muppidi et al., 2015). Our data show for the first time that an FDC-derived metabolite can be locally maintained and act directly on GC B cells. Other cells have been reported to express *Ch25h*, but anatomy places FDCs at a prime location to directly control GC B cells via 25-HC in an EBI2 independent fashion, without requiring long range tissue diffusion.

Moreover, we demonstrated that increasing cholesterol in the diet is sufficient to modulate 25-HC tissue concentration in PPs and to reduce Ag-specific IgA response. Our data do not exclude that in the presence of increased concentration of tissue cholesterol, cells other than FDCs could produce 25-HC outside of the GC, and therefore contribute to PC inhibition at different stages of B cell activation.

25-HC’s ability to block activation, and subsequent transcriptional activity of SREBP2, an ER protein that senses intracellular sterol concentration, is well documented, but its regulation remains largely unexplored in adaptive immune cells. B cells activated *in vitro* by IgM BCR stimulation upregulate SREBP2. Similarly, LZ GC B cells, that routinely test their BCR against Ag displayed by FDCs, are also characterized by increased SREBP2 induction and transcriptional activity. We suggest a model in which SREBP2 activation is modulated by 25-HC in the LZ to reduce PC differentiation. Alterations of SREBP2 levels sharply affect GC B cell fate, suggesting that environmental cues able to control SREBP2 activation might play a role in GC output in addition to the widely studied BCR affinity and T cell help.

Although our over-expression results suggested a striking effect for SREBP2 activation in PC differentiation, the *in vivo* phenotype in mice lacking the SREBP2 inhibitor 25-HC was only evident when Ag-specific response was tracked and was reduced in magnitude. While our data suggest a dominant role for CH25H in mediating dietary cholesterol inhibition of PC response, we cannot exclude the possibility that other oxysterols or cholesterol itself could play a compensatory role in absence of 25-HC. In line with the negative feedback exerted by cholesterol and selected oxysterols on cholesterol metabolism via SREBP2 inhibition, albeit at higher concentration, CH25H-independent PC inhibition could take place when cholesterologenic diet is prolonged or contains an even higher amount of cholesterol.

Moreover, external cues could modulate the strictness of 25-HC/SREBP2 crosstalk in GC B cells, including diet and commensal composition. The original report describing *Ch25h*^*-/-*^ mice reported increased IgA production (Gold et al., 2014): while the authors explained it as an effect on CSR based on *in vitro* experiments, we could not see 25-HC inhibition of IgA CSR in PP or MLN in our mouse colony. However, the initial IgA phenotype reported by Russell could be easily reconciled with our data by invoking a SREBP2 activation and increased PC differentiation mediated by a stronger T cell dependent commensal response in their colony.

We propose a model in which FDCs convert dietary cholesterol into 25-HC in PPs which diminishes Ag-specific PC responses. Given that cholesterol absorption is dependent on bile acid levels, and that selected members of the microbiome can degrade bile acid, we posit that early dysbiosis will result in decreased dietary sterol absorption and consequent 25-HC levels. Since multiple antigens simultaneously drive chronic intestinal GCs, a metabolite-based mechanism to tilt the GC outcome might provide a rapid mode of Ab regulation in response to fluctuations in the luminal commensal composition. SREBP2 has a well-established role in controlling cholesterol metabolism in non-hematopoietic cells: our findings here have established a critical role for SREBP2 in controlling GC B cell fate. While *in vitro* differentiation into PCs was inhibited in absence of SREBP2, the nature of the genetic model we used prevented us from uncovering the intrinsic role of SREBP2 in PC survival. Our *in vitro* data suggest that BCR stimulation is the dominant signal for SREBP2 transcription and activation and it is likely to shape GC B cell fate in the LZ; however, we cannot rule out a role for T cells in maintaining SREBP2 activation and promoting differentiation in PCs.

The SREBP2 signature can be clearly observed in LZ GCs of mouse (Radtke and Bannard, 2019) and human (Holmes et al., 2020), suggesting an evolutionary conserved mechanism to turn on cholesterologenic genes upon selection in the LZ. Therefore, our findings regarding metabolic requirements for controlling PC differentiation may be of broader relevance to understand humoral response in human populations exposed to high cholesterol diet.

## Materials and Methods

### Mice

C57BL/6J (CD45.2) (Stock No: 00064), Ly5.2 (CD45.1) congenic C57BL/6 (B6) (Stock No 002014), Rosa26DTR (Stock No: 007900), Srebp2 flox/flox (Stock No: 031792), *Ch25h*^-/-^ (Stock No: 016263), *Gpr183*^-/-^ (Pereira et al., 2009) AID cre (Stock No: 007770), CD21^cre^ (Stock No: 006368), TdTomato (Stock No: 007914) were purchased from the Jackson Laboratory. All mice were bred and maintained under standard 12:12 hours light/dark conditions and housed in specific pathogen-free (SPF) conditions at the University of Massachusetts Medical School. All procedures were conformed to ethical principles and guidelines approved by the UMMS Institutional Animal Care and Use Committee.

### Mouse Diets

Mice were either fed a standard chow diet (ISO-PRO 3000 sterile rodent diet #5P76 (LabDiet)) for the duration of the experiment or a high cholesterol diet where 0.15% cholesterol is added to the ProLab RMH 3000 5P76 diet. (0.15% HCD Envigo TD.180381customized diet).

### Immunizations, infections, and treatment

For cholera toxin responses, mice were immunized with 10 ug of cholera toxin (List Biological Laboratories) in PBS by oral gavage. Animals received cholera toxin every 7 days for three consecutive weeks. Mice were analyzed 7 days after last immunization. Metabolically defective (aroA) *S. typhimurium* strain SL1344 was provided by Milena Bogunovic lab (UMMS) and was grown at 37°C in Luria broth supplemented with appropriate antibiotics to preserve mutation and plasmid. Mice were orally gavaged three times on alternate days with 10^9^ CFUs of aroA S. typhimurium in 200 ul 5% sodium bicarbonate. Serial dilutions of bacterial preparations were plated onto LB-agar plates to confirm administered dose.

To prevent lymphocyte egress from Peyer’s patches, the S1PR1 agonist FTY-720 (Fingolimod-HCl Cat No.S5002 from Selleck Chemicals) was dissolved in saline solution and administered to mice daily at 1mg/kg for 3-7 days as detailed in the figures.

### Cell Isolation

Peyer’s patches and mesenteric lymph nodes were digested in digestion media (RPMI, 5% heat inactivated FBS, 10 mM HEPES, 1% P/S, 50 ug/mL DNase I (Sigma DN25), and 0.5 mg/mL Collagenase IV (Worthington Biochemical LS004189)) rotating at 37°C for 15 min. The digested tissue was then smashed through 70μm cell strainer. Spleens were harvested and smashed through 70μm cell strainer. Isolated cells were then washed with FACS buffer (1X DPBS, 2% heat inactivated FBS, 2mM EDTA) and counted for further analysis. For isolation of lamina propria (LP) lymphocytes, small intestine was dissected and flushed with cold PBS and Peyer’s Patches were removed. The small intestine was divided into 3 equal parts, the proximal (duodenum) and distal (ileum) were opened longitudinally and vortexed in a 50ml conical tube containing HBSS supplemented with 5% heat-inactivated FBS and 10mM Hepes. Epithelial cells were removed by rotating the small intestine tissue in pre-digestion media (RPMI medium, 5% heatinactivated FBS, 10mM Hepes, 10mM EDTA) for 30 minutes at 37°C. The intestinal pieces were then washed with complete media (10% heat inactivated FBS, 10 mM HEPES, 1% P/S), chopped with scissors, and digested at 37°C for 30 minutes in digestion media. Digested tissue was passed through 70 um cell strainer and isolated cells were resuspended in 40% Percoll-RPMI and layered with 80% Percoll-RPMI and subsequently centrifuged for 20min at 2200 rpm without break. The isolated LP cells were enumerated on a BD LSRII using AccuCheck Counting Beads (Invitrogen) as per manufacturer recommendations.

### Flow cytometry and cell sorting

Mesenteric lymph node, Peyer’s patches, and spleen were collected as described above. Cell suspensions were stained with LIVE/DEAD Fixable Aqua Dead Cell Stain (Invitrogen) or Fixable Viability Dye eFluor780 (Invitrogen 65086514) in FACS buffer, and Fc receptors blocked with anti-mouse CD16/32 (2.4G2). BD Cytofix/Cytoperm kit (BD BDB554714) was used for fixation and intracellular staining. Cells were incubated for 20 min on ice with antibodies to B220, IgA, IgD, IgM, IgG1, IgG2b, IgG3, GL7, CD138, CD38, CD45.1, CD90.2, CXCR4 and CD86 (the antibodies used are described in table S1). Data were collected on a BD LSR II and analyzed in FlowJo v10.7 software.

For cell sorting, cells were stained as described above and sorted on a BD FACSAria II with a 70 or 84 micron nozzle. Cells were maintained at 4°C until sorting. Sorted plasma cells from PP and mLN were used for either ELISPOT assay or analysis. Germinal center, Follicular B cells, Memory and plasma cells were sorted as live IgD-GL7+ CD38-CD138-, live IgD+ GL7-CD138-, live IgD-GL7+ CD38+ CD138-, IgD-GL7-CD38-CD138+ respectively into TRIZOL (Invitrogen) for RNA extraction or into tissue culture plates containing complete media for further analysis. Light zone and dark zone germinal center B cells were sorted based on expression of CXCR4 and CD86.

### RNA extraction and Real Time PCR

Total RNA was isolated with 0.5 mL of TRIZOL reagent (Invitrogen) following the manufacturer’s protocol. Reverse transcription was performed using Superscript III reagent kit from Invitrogen. All samples were checked for quality (A260/A280 ratio of 1.8-2.0). Real Time PCR was performed by using iQ SYBR Green superMix (BioRad). Gene expression levels were determined using a comparative method (DCq) normalizing the target mRNA to b actin as endogenous internal control. Forward and reverse primers sequences are listed in table S2.

### Enzyme-linked ImmunoSorbent Assay (ELISA)

Ninety-six–well half area high binding flat bottom plates (Corning) were coated with 25ul of 2 μg/ml purified anti-IgA (RMA-1, BD), CT (List biological laboratories) diluted in PBS overnight at 4°C. Plates were washed and blocked with PBS–5% BSA before diluted intestinal wash or fecal samples were added and threefold serial dilutions were made. Samples were processed as described previously by *Reboldi et al*. and incubated overnight at 4°C. Bound antibodies were detected by anti-IgA-conjugated horseradish peroxidase (Southern Biotech) and visualized by the addition of Substrate Reagent Pack (Biolegend). Color development was stopped with 3 M H2SO4 stop solution (Biolegend). Purified mouse IgA (Southern Biotech) served as standard. Absorbances at 450 nm were measured on a tunable microplate reader (VersaMax, Molecular Devices). Antibody titers were calculated by extrapolating absorbance values from standard curves where known concentrations were plotted against absorbance using SoftMax Pro 5 software. OD was plotted for CT due to lack of standard curve.

### ELISpot

ELISpot plates (Millipore) were coated with 100μl of 2 μg/ml purified anti-IgA in PBS overnight at 4°C. Plates were washed three times with PBS then blocked for 2 h at 37°C with 10% FCS-RPMI. Cells were isolated from Peyer’s patches, mesenteric lymph nodes and lamina propria and counted as described above then diluted in blocked ELISpot plates and incubated overnight in a 37°C 5% CO2 tissue culture incubator. The next day, plates were washed three times with PBS-0.1% Tween then PBS.

Detection of IgA spots was achieved by using anti-IgA-conjugated horseradish peroxidase and developed with 3-amino-9ethylcarbazole (Sigma-Aldrich). Color development was stopped by washing several times with water. Once dried, plates were scanned and spots counted using the CTL ELISPOT reader system (Cellular Technology).

### NB-21 coculture

Germinal center B cells from Peyer’s patches and mesenteric lymph nodes were FACS sorted and cocultured with NB-21 feeder cells as previously described and with adjustments (Kuraoka et al., 2016; Stewart 2018). Briefly, 10^3^ NB-21.2D9 cells/well in 100 uL of B cell media (RPMI supplemented with 10% heat-inactivated FBS, 10 mM HEPES, 1mM sodium pyruvate, 100 units/mL penicillin, 100 mg/mL streptomycin, MEM nonessential amino acid and 55 mM 2-Mercaptoethanol) were seeded into 96-well plates 1 day before B cell co-culturing. On the next day, 10^4^ B cells in 100 uL of B cell media was added to each well. 100uL of media was replaced on day 2. Cholesterol, 7a,25-HC or 25-HC were added to the B cell/NB-21 co-culture at days 0 and 2. Culture plates were centrifuged at day 3.5 and supernatant was collected for IgA detection.

### Nuclei Preparation

Nuclei preparations were carried out as previously described (Gallagher, M. P Immunohorizons, 2018). Briefly, follicular B cells were isolated from spleens of wild type or *Aicda*^*cre/+*^ *Srebf2*^*fl/fl*^ mice using EasySep mouse B cell isolation kit and protocol (StemCell). Isolated cells were then cultured *in vitro* in the presence or absence of 5 mg/ml anti-IgM F(ab)_2_ (Jackson Immunoresearch) and/or 5 mg/ml anti-CD40 (BioXCell) for 16 or 48 hours. Inhibition experiments were carried out by culturing isolated follicular B cells with 5 ug/ml anti-IgM in the presence or absences of 10 nM Ibrutinib or 50 µM Rapamycin (Selleck Chemical) for 16 hours. Following the culture, the cells were harvested and lysed in Sucrose Buffer A (10 mM HEPES (Gibco), 8 mM MgCl_2_, 320 mM Sucrose, 0.1 % (v/v) Triton-X 100 (Sigma), and 1X complete, EDTA-free Protease Inhibitor Cocktail (Roche) on ice for 15 minutes. The cells were then centrifuged at 2000g for 10 minutes and washed twice with Sucrose Buffer B (Sucrose Buffer A without Triton-X 100). Nuclei were fixed in 4% Paraformaldehyde (electron microscopy grade, Electron Microscopy Sciences), diluted in Sucrose Buffer B, at room temperature for 25 minutes. Nuclei were stained as described in figure 4 and figureS4 and analyzed by flow cytometry.

### Bone marrow chimeras

CD21^cre/Rosa26DTR+^or CD21^cre^ mice were lethally irradiated twice with 550 rads gammairradiation, 3 hours apart, then intravenously injected with 1-3 × 10^6^ bone marrow cells. Bone marrow was harvested by flushing both tibia and femurs of donor mice. Mice were analyzed 8-12 weeks later.

### FDC ablation

For LT-blocking experiments, animals were treated with LTβR-Fc using 100 μg on days −3 and −1 before sacking the mice. FDC ablation with DT was performed as described previously (Wang et al., 2011). In brief, animals expressing CD21-cre and ROSA-DTR alleles or littermate controls were lethally irradiated and reconstituted with WT BM. 6–8 wk after reconstitution, animals were treated with 100 ng DTx (EMD Biosciences) i.p. and analyzed 16–20 h later.

### Lipid extraction and 25HC/7a,25-HC quantification

Lipids from 100 mg of Peyer’s patches and mesenteric lymph nodes were extracted using the Folch method for lipid extraction (Folch et al., 1957). Lyophilized lipid was dissolved at 100 mg/ml in ethanol. 25-HC was measured by using 293T cells supernatant from cells transfected sequentially with *pENTR-Hsd3b7* and *pENTR-Cyp7b1* vectors. The 7a,25-HC activity was evaluated by transwell chemotaxis assay. Lipid extracts were diluted in 10 volumes of sterile chemotaxis media (RPMI + 0.5% fatty acid free BSA) and tested for EBI2 dependent bioactivity by transwell chemotaxis assays of 50:50 mixed M12 B cell line transduced with an EBI2-IRES-GFP retroviral construct and mock M12 cells. The three-hour migration assay was performed at 37°C and migrated cells were stained for DAPI and analyzed by flow cytometry. The relative migration of EBI2-GFP+ M12 cells over M12 cells was normalized to migration toward lipid free migration media. The purified 7a,25-HC was used as positive control for chemotaxis.

### Statistical analysis and quantification

Statistical analyses were performed using GraphPad Prism v8.0 using two-way ANOVA with Bonferroni’s multiple comparisons test or unpaired T test Data are presented as means +/-SEM. Differences between group means were considered significant at indicated p value. Sample sizes can be found in the figure legends, indicating the number of animals or cells used for each experiment.

## Author Contributions

AR conceived of the project and experiments. BCT, SC, AB, FR, KH, JRM, and AR performed experiments and analyze the data. AR wrote the initial manuscript. All authors contributed to manuscript review.

## Acknowledgements

This work was supported by MMRF Research Fellowship and LLS New Idea Award (A.R.). We thank Coral Freeman for mouse husbandry.

---

**Table S1.**

| **Antibody** | **Clone** | **Vender** |
| --- | --- | --- |
| *Ultra-LEAF™ Purified anti-mouse CD16/32* | 93 | Biolegend |
| *anti-mouse/human CD45R/B220* | RA3-6B2 | Biolegend |
| *IgA Rat anti-Mouse* | C10-1 | BD |
| *anti-mouse IgD* | 11-26c.2a | Biolegend |
| *anti-mouse IgM* | RMM-1 | Biolegend |
| *anti-mouse IgG1* | RMG1-1 | Biolegend |
| *anti-mouse IgG2b* | RMG2b-1 | Biolegend |
| *anti-mouse IgG3* | RMG3-1 | Biolegend |
| *anti-mouse/human GL7 Antigen* | GL7 | Biolegend |
| *anti-mouse CD138 (Syndecan-1)* | 281-2 | Biolegend |
| *anti-mouse CD38* | 90 | Biolegend |
| *anti-mouse CD45.1* | A20 | Biolegend |
| *anti-mouse CD90.2* | 30-H12 | Biolegend |
| *anti-human CD184 (CXCR4)* | 12G5 | Biolegend |
| *anti-rat CD86* | 24F | Biolegend |
| *SREBP2 Polyclonal Antibody (a.a. 843-857)* |  | ThermoFisher Scientific |
| *anti-SREBP-2 Antibody* | 22D5 | MilliPORE SIGMA |
| *SREBP2 Polyclonal Antibody (a.a. 1-220)* |  | ThermoFisher Scientific |

**Table S2.**
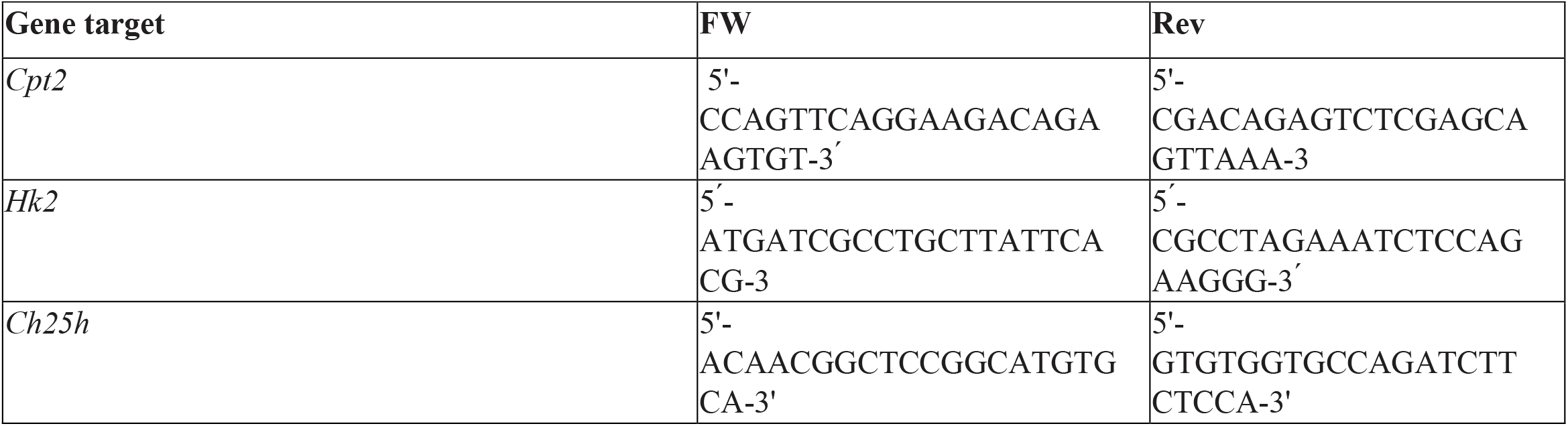
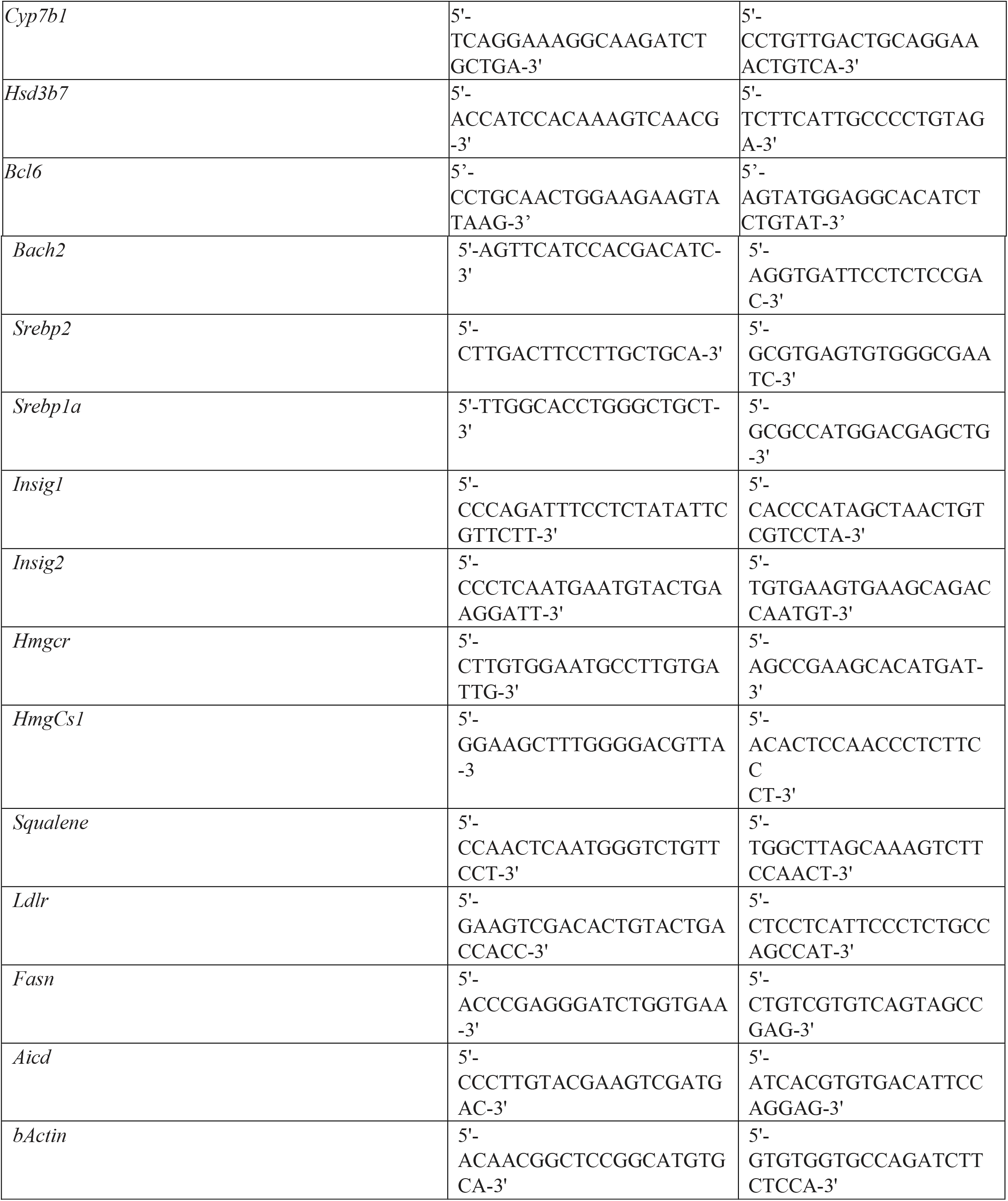

