## Supplemental figures for "Intracellular sterol sensing controls intestinal B cell differentiation" (from another version of this preprint: version 2, 2020-12-17)

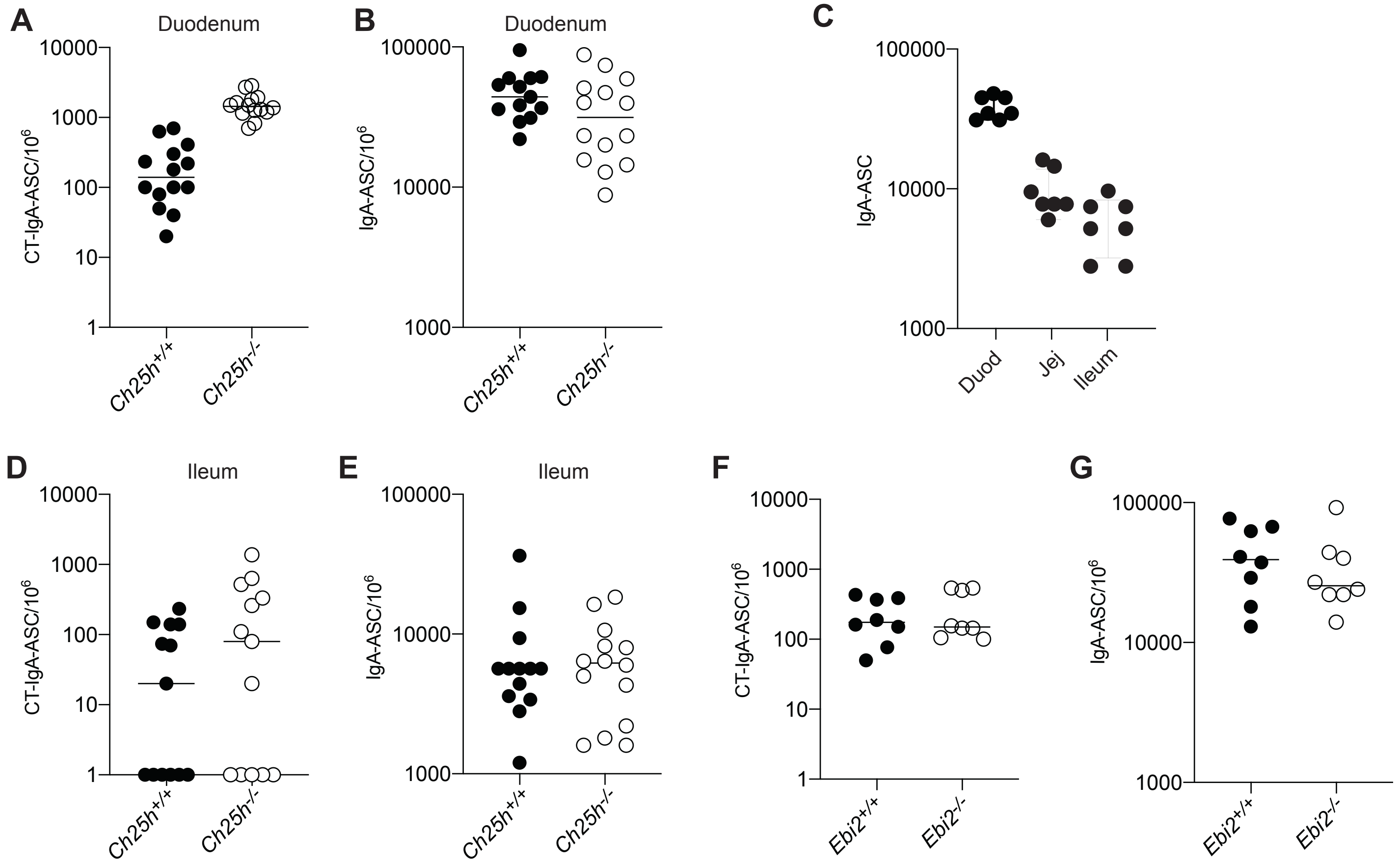


**Supplementary Figure 1**

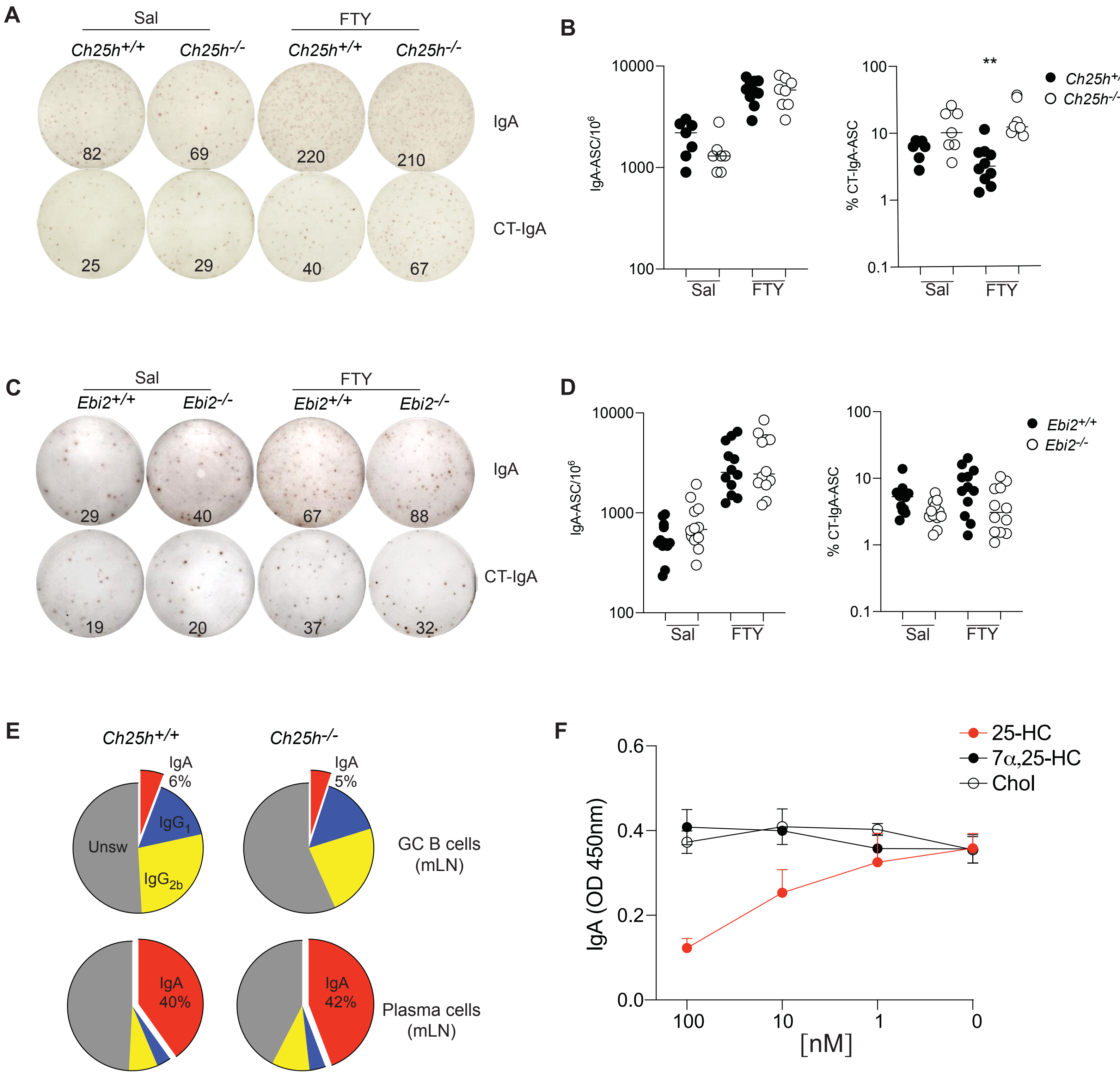


**Supplementary Figure 2**

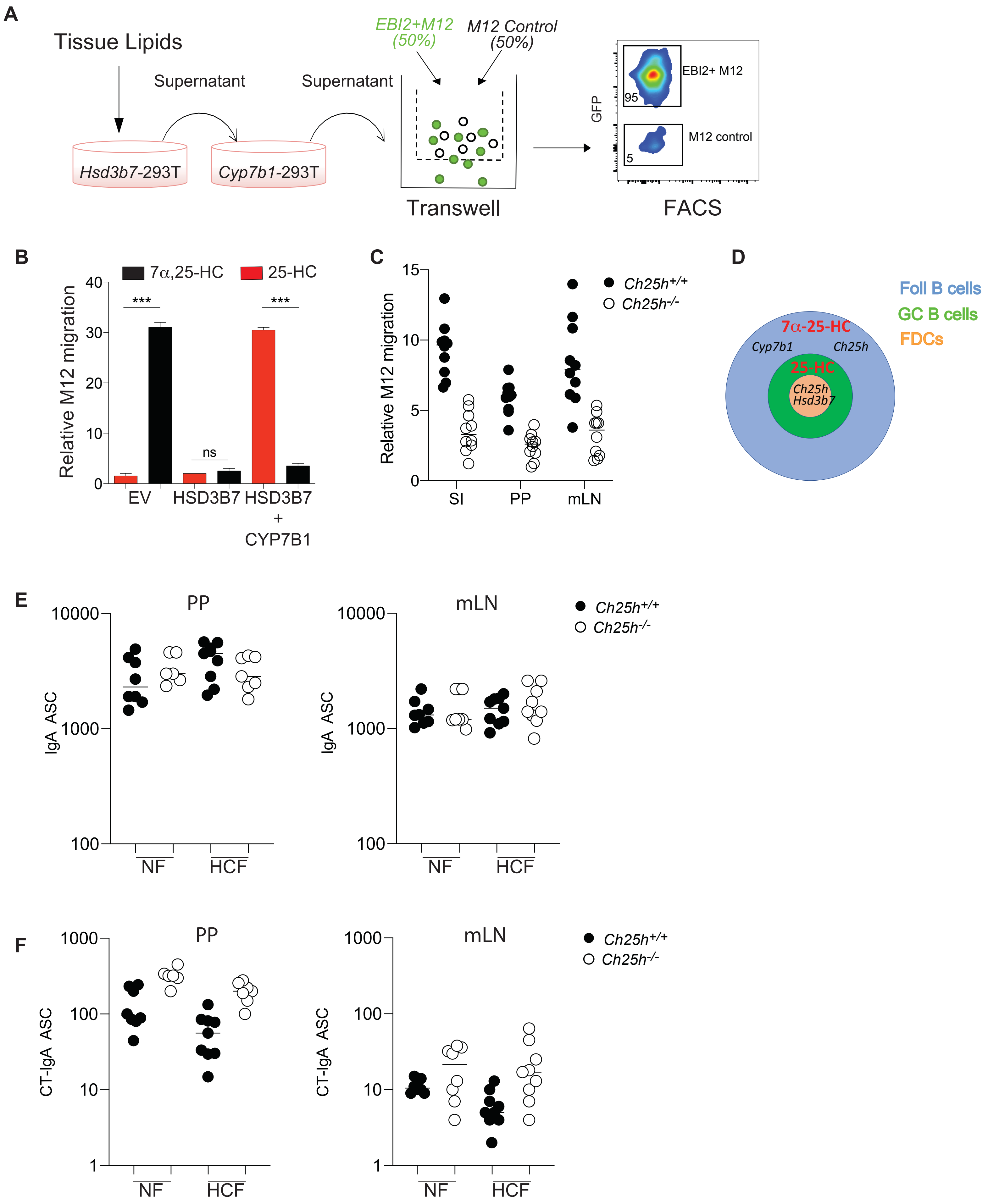


**Supplementary Figure 3**

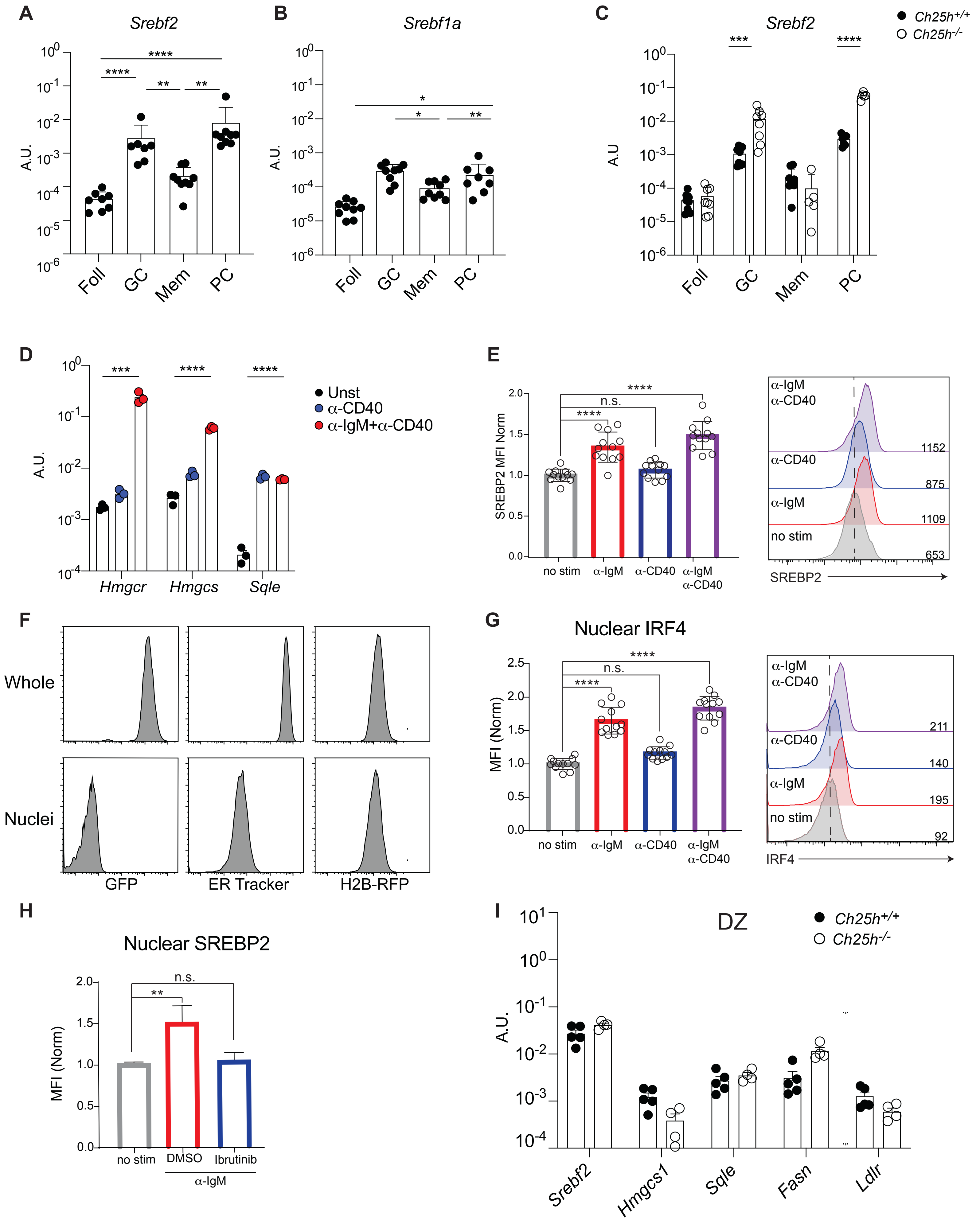


**Supplementary Figure 4**

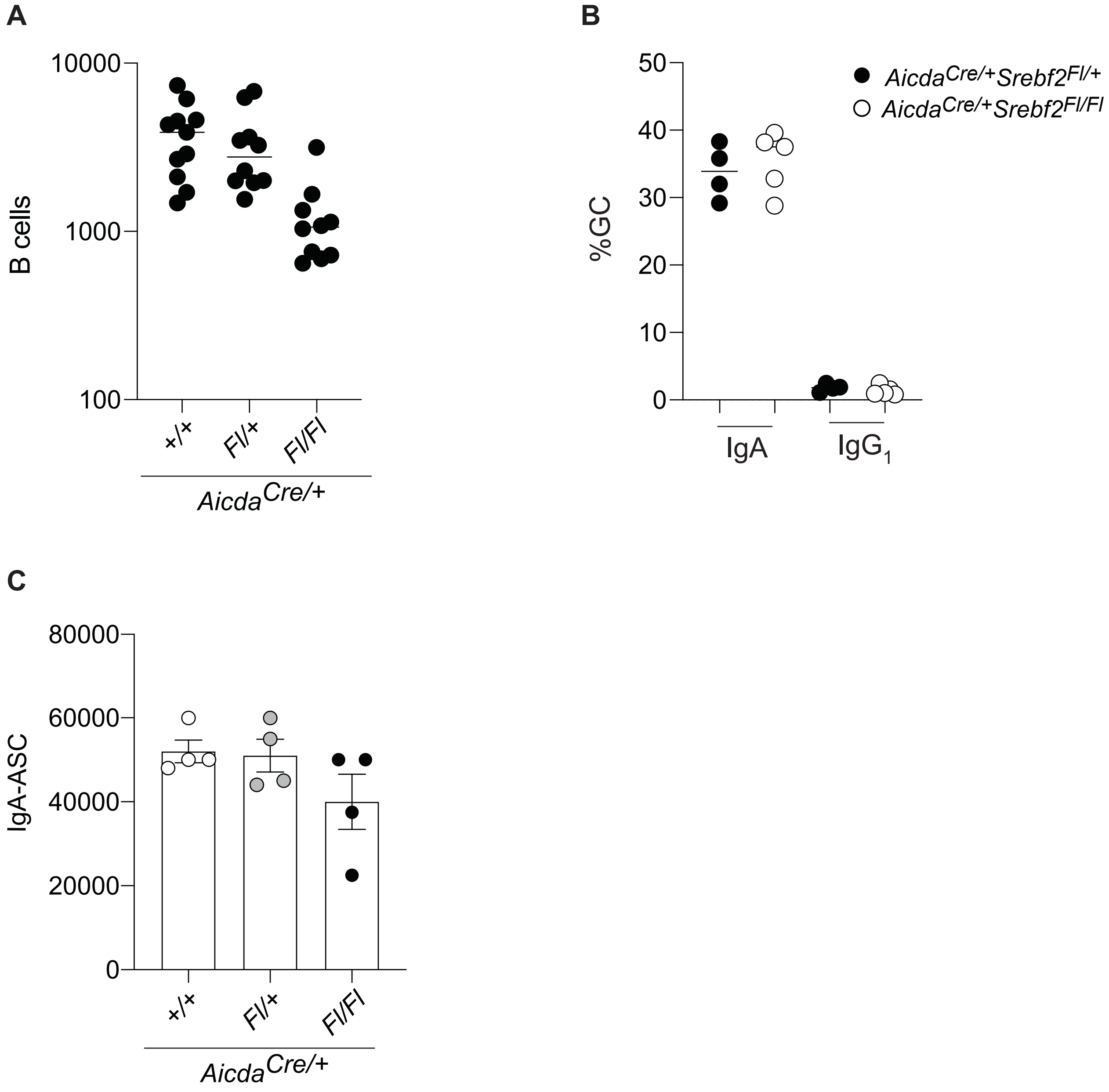


**Supplementary Figure 5**

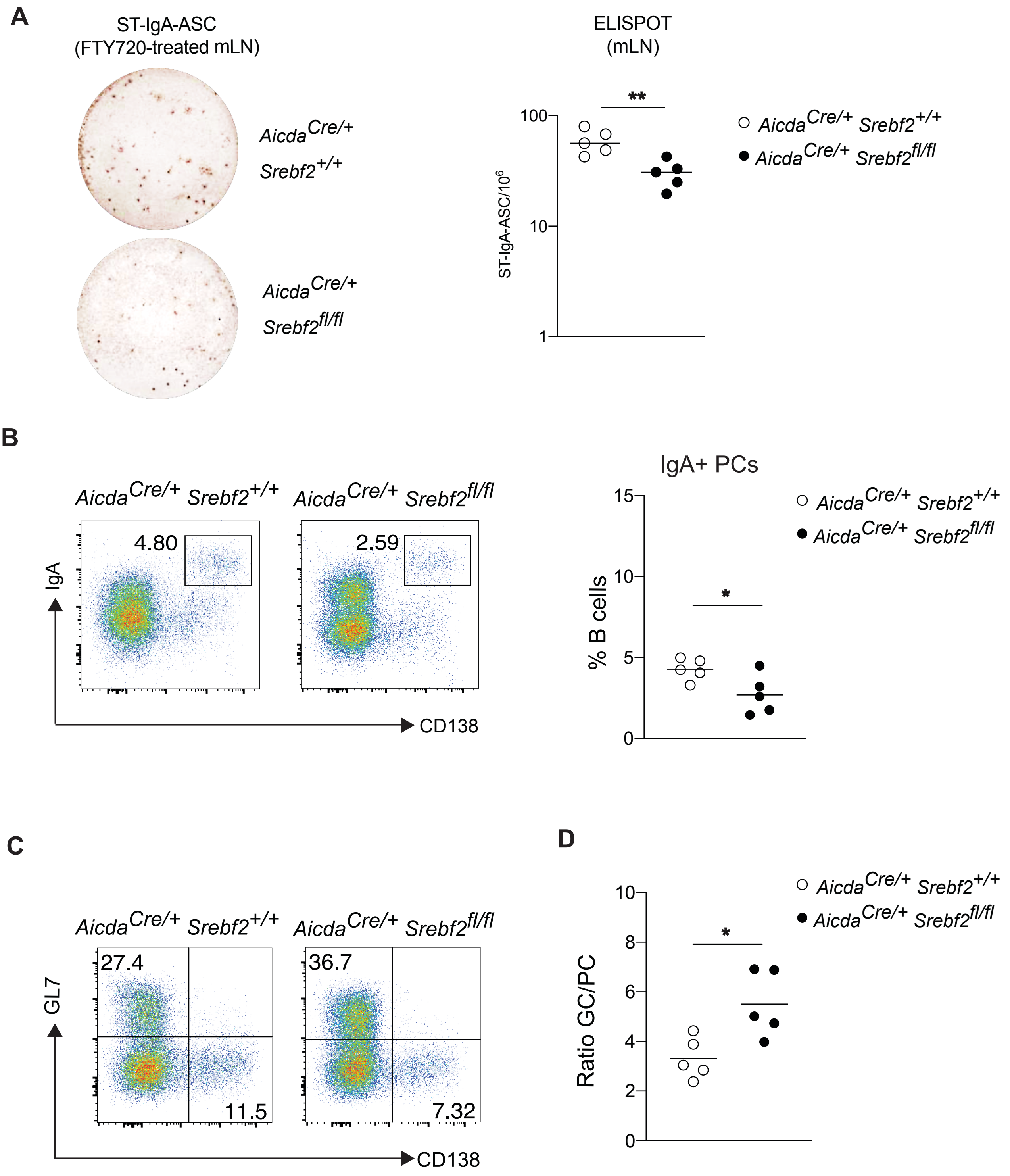


**Supplementary Figure 6**

### Supplementary figure legends

**Figure S1. 25-HC controls antigen-specific IgA secretion in duodenum, but not in the ileum.**
(A) ELISPOT of CT-specific IgA and (B) total IgA secretion in duodenum of *Ch25h*$^{-/-}$ and LMC mice. (C) Number of secreting IgA PC in Duodenum, Jejunum and Ileum, analyzed by ELISPOT. (D) CT-specific IgA ASC and (E) total IgA in the ileum of *Ch25h*$^{-/-}$ and LMC mice, depicted by ELISPOT. (F,G ) ELISPOT data showing IgA ASC and total IgA in duodenum of *Ebi2*$^{-/-}$ and LMC mice Graphs show data from $3^3$ experiments. *p<0.05, **p<0.01,***p<0.005 (ANOVA multiple comparation).

**Figure S2. 25-HC modulates CT-specific IgA response in PPs independently of class switching.**
(A) Image of representative ELISPOT of PP IgA and CT-specific IgA secreting cells from *Ch25h*$^{-/-}$ and LMC mice orally immunized with CT for 3 weeks and treated for one week with FTY720 or saline by IP injection.(B) Graphs showing the plotted analysis of the total number of IgA secreting PC and the frequency of the CT-specific IgA PC, detected by ELISPOT assay depicted in (A).(C) Representative ELISPOT of PP IgA and CT-specific IgA secreting cells from *Ebi2*$^{-/-}$ and LMC mice treated for 3 weeks with CT by oral gavage and one week with FTY720 or saline by IP injection. (D) Total IgA and CT-specific IgA ASC in PPs of *Ebi2*$^{-/-}$ and LMC mice. (E) Class switch recombination in GC B cells and PC from mLN of the indicated genotype assessed by FACS. (F) Quantification of secreted IgA by ELISA from GC B cells cultured with NB21 supernatant and anti-CD40 and stimulated with the indicated sterols for 3.5 days.
Graphs in B,D and F depict the mean from 2-3 independent experiments *p<0.05, **p<0.01,***p<0.005 (two-way ANOVA)

**Figure S3. 25-HC quantification in PPs and antigen-specific IgA in response to dietary cholesterol.**
(A) Schematic illustration of transwell migration assay with EBI2$^{+}$ M12 for the quantification of 25-HC in intestinal SLO using transfected 293T cell. (B) Validation of the strategy in (A) using synthetic 25-HC and 7a,25-HC. Graph shows the relative migration of EBI2$^{+}$ M12 cells from 3 experiments. (C) 25-HC quantification in the indicated tissue from *Ch25h*$^{-/-}$ and LMC mice. (D) Model for 25-HC rich niche in GC. (E) ELISPOT of IgA and (F) CT-specific secreting cells in PPs and mLN of mice treated for 1 week with regular chow or 0.15% cholesterol supplement diet. ELISPOT data were pooled from at least 3 independent experiment *** p<0.005 (unpaired Student's t test (B) and two-way ANOVA (D, E)).

**Figure S4. 25-HC reduces SREBP2 and alters SREBP2-mediated transcription in vitro and in vivo.**
Transcript level of *Srebf2* (A) and *Srebf1a* (B) in B cell subsets from mLN of wild-type mice and (C) from mLN of *Ch25h*$^{/-}$ and LMC mice measured by qPCR. (D) qPCR of mevalonate pathway and SREBP2 targets on B cells stimulated for 48hrs with aCD40 or with aCD40 with aIgM. (E) Mean Fluorescence index (MFI) and representative histograms of total SREBP2 detected by flow cytometry in B cells stimulated for 16h with the indicated stimuli. (F) Whole cell and nuclei isolation stain of transduced WEHI for presence of cytoplasmic, ER and nuclear marker. (G) Mean Fluorescence index (MFI) of intranuclear IRF4 detected by flow cytometry in B cells stimulated for 16h with the indicated stimuli. (H) Mean Fluorescence index (MFI) of nuclear SREBP2 detected by flow cytometry in B cells stimulated for 16h with aIgM and the indicated inhibitors. (I) Gene expression of *Srebf2* and SREBP2- target genes in sorted DZ B cells (gated on CXCR4+ CD86- cells ) from mLN of *Ch25h*$^{-/-}$ and LMC mice. Data were produced from at least experiment 2-3 independent experiment. *p<0.05, **p<0.01,***p<0.005, **** p<0.001 (two-way ANOVA)

**Figure S5. SREBP2 deficiency impacts GC B cell differentiation but not IgA class switching.**
(A) Number of B cells from the indicated genotype recovered after 2.5 days of NB21 culture. (B) IgA and IgG1 class switch in GC B cells from the indicated genotype. (C) ELISPOT of IgA secreting PC in lamina propria of the mice of the indicated genotype.

**Figure S6: IgA response to enteric infection in mLN is controlled by SREBP2 in GC B cells.**
(A) Representative ELISPOT and compiled data showing Salmonella-specific IgA secreting cells in mLN of *Aicda*$^{cre}$ *Srebf2* $^{+/+}$ and *Aicda*$^{cre}$ *Srebf2* $^{flox/flox}$ mice infected with Salmonella and treated with FTY720. (B) Representative flow cytometry and frequency of IgA+ PCs from mLN of *Aicda*$^{cre}$ *Srebf2* $^{+/+}$ and *Aicda*$^{cre}$ *Srebf2* $^{flox/flox}$ mice infected with Salmonella and treated with FTY720 as in A. (C) Representative flow cytometry of GC B cell and PC from mLN of *Aicda*$^{cre}$ *Srebf2* $^{+/+}$ and *Aicda*$^{cre}$ *Srebf2* $^{flox/flox}$ mice infected with Salmonella and treated with FTY720 as in A, B. (D) Ratio of GC B cell and PC

from PPs from C. Each symbol represents one mouse from 2 independent experiments. *p<0.05, **p<0.01 (two-way ANOVA).
